# The response regulator BqsR/CarR controls ferrous iron (Fe^2+^) acquisition in *Pseudomonas aeruginosa*

**DOI:** 10.1101/2025.04.12.648518

**Authors:** Alexander Paredes, Mackenzie Hull, Harvinder Singh, Darryn Greene, Ahmed O. Tajudeen, Aya Kubo, Sara Patamawenu, Rachel Kramer, Janae Baptiste Brown, Kelly N. Chacón, Marianna A. Patrauchan, Aaron T. Smith

**Author notes:** These authors contributed equally to this work (Alexander Paredes, Mackenzie Hull).

## Abstract

*Pseudomonas aeruginosa* is a ubiquitous, Gram-negative bacterium that forms biofilms and is responsible for antibiotic-resistant hospital-acquired infections in humans. The *P. aeruginosa* BqsRS two-component system regulates biofilm formation and dispersal by sensing extracytoplasmic Fe^2+^, but the mechanistic details of this process are poorly understood. In this work, we report the crystal and solution structures of the *Pa*BqsR response regulator receiver domain, comprising a (βα)_5_ response regulator assembly, and the DNA-binding domain, comprising a helix-turn-helix motif. Consistent with its cognate stimulus being Fe^2+^, we show that *Pa*BqsR binds directly to the promoter region of the *feo* operon that encodes the bacterial Fe^2+^ transport system FeoABC. Corroborating these *in vitro* results, transcriptional studies show that *Pa*BqsR is a global regulator controlling many important genes in PAO1, including the *feo* operon. Intriguingly, promoter-based assays reveal that *Pa*BqsR is a dynamic regulator that responds to bioavailable Fe^2+^, likely through the ability of *Pa*BqsR to bind Fe^2+^ directly via a His-rich motif, independent of the *Pa*BqsS membrane His kinase. This mode of regulation is unprecedented among OmpR-like response regulators but represents an important level of control over Fe^2+^ acquisition in *P. aeruginosa* that could be an attractive therapeutic target to treat hospital-acquired infections.

## INTRODUCTION

Bacteria are robust microorganisms that are capable of surviving in numerous, diverse conditions due to their abilities to sense, respond, and adapt to dynamic environments.^1–5^ Most bacteria utilize two-component signal transduction systems (TCSs) to mediate communication between extracellular and intracellular milieus in order to facilitate this adaptation.^6,7^ TCSs are widespread among bacteria and regulate critical cellular processes like motility, chemotaxis, the production of virulence factors, and the acquisition of essential nutrients such as transition metals.^1–8^ The most common type of TCSs are membrane-bound and canonically combine two proteins to serve three functions: environmental recognition, signal transduction, and (most commonly) transcriptional regulation (Fig. 1). The two typical proteins comprising a TCS are the transmembrane sensor His kinase (HK) and the cytoplasmic response regulator (RR) (Fig. 1). The membrane-bound sensor HK receives the environmental stimulus by interacting directly with a signaling ligand, thus beginning the signal transduction process. Stimulus binding then induces HK autophosphorylation (HK-P_i_),^9^ after which the phosphate signal may be readily transferred to a conserved residue on the RR (RR-P_i_) (Fig. 1).^9–11^ Phosphorylation of the RR occurs at the N-terminal regulatory domain (site of a conserved Asp residue), triggering conformational and dynamical changes in the protein, commonly altering the affinity of the RR to DNA and affecting transcriptional regulation (Fig. 1).^9, 12^ Stimulus-specific HKs are generally paired with cognate RRs that activate or (less frequently) inhibit transcription of DNA when bound, thus regulating bacterial responses to external inputs.^9, 13–14^ Since these systems facilitate the ability of bacteria to adapt to diverse environments, TCSs also function as critical virulence factors that enhance the infections of pathogens within hosts.^15–18^

**Figure 1.**
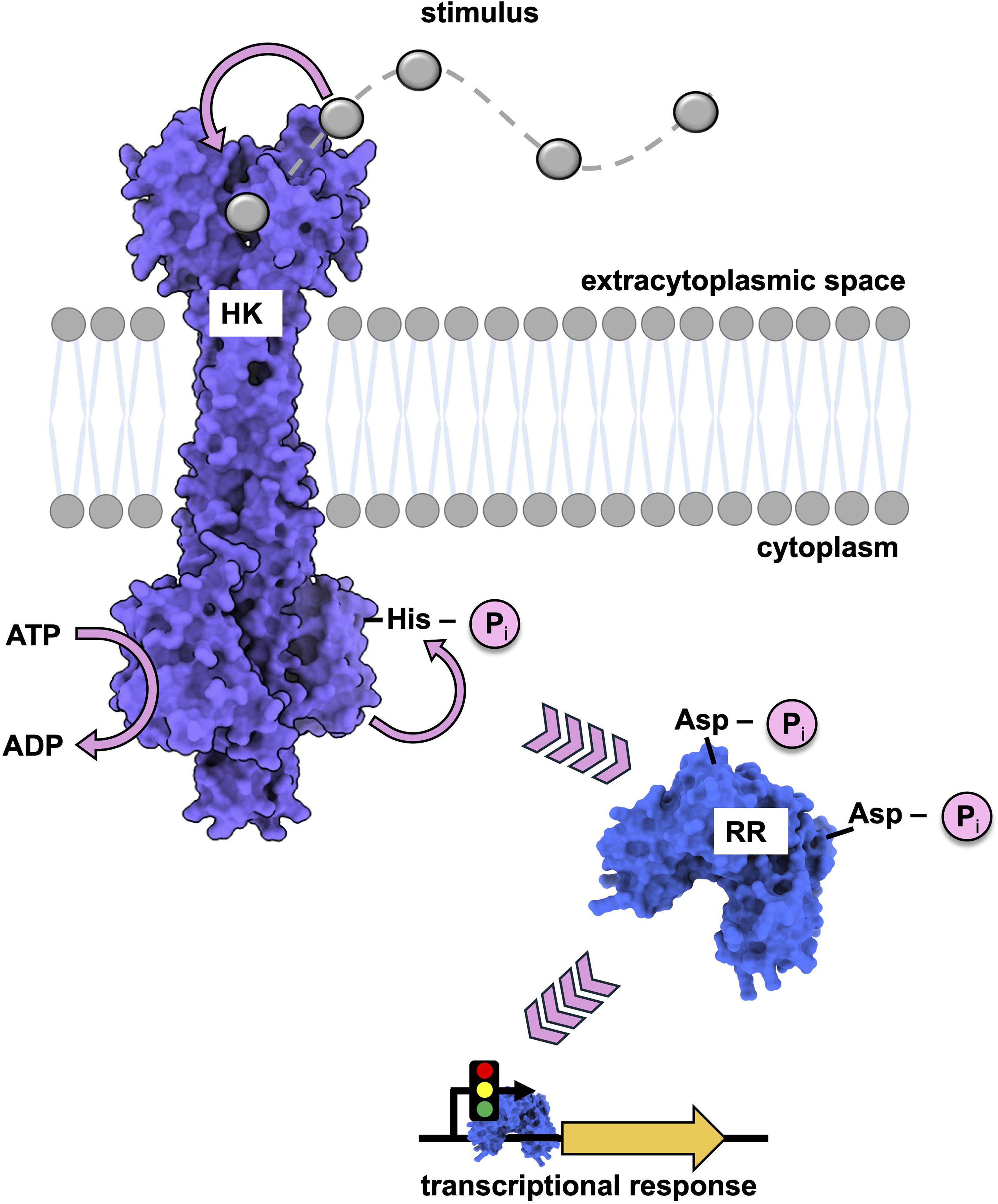
The signal transduction pathway of a canonical two-component system (TCS). Membrane-bound His kinases (HK) generally function as homodimers; when activated by an extracytoplasmic signal, the cytosolic kinase catalytic core of one HK monomer phosphorylates the same subunit of the neighboring dimer by transferring the γ-phosphate of ATP to the conserved His residue located in the dimerization and His phosphotransfer (DHp) domain. Upon the completion of this autophosphorylation event, the linked cytosolic response regulator (RR) accepts the transfer of the phosphate group to a conserved Asp within its N-terminal regulatory domain. As a result, the phosphorylated RR undergoes dynamical and conformational changes (generally dimerization) that modify the affinity of the RR to particular genes, ultimately leading to an altered transcriptional response.

One increasingly problematic pathogen that makes extensive use of TCSs to adapt to its host is *Pseudomonas aeruginosa*, a ubiquitous and opportunistic Gram-negative bacterium that can reside within virtually all environments and is most notably associated with hospital-acquired infections.^2,12^ As one of the six multidrug-resistant ESKAPE pathogens,^13–15^ *P. aeruginosa* continues to be a global health threat and is well-known for infecting human patients such as those with burn wounds, neutropenia, and/or those suffering from cystic fibrosis (CF).^12^ As part of its outstanding adaptability, *P. aeruginosa* can form biofilms, protective and intricate three-dimensional aggregates composed of polysaccharides, proteins, lipids, and DNA.^16–18^ Bacteria within these complex structures restrict antibiotic penetration, have increased horizontal gene transfer of antibiotic resistance genes, and are protected by the physical defenses of the host.^16–18^ Biofilms enhance the ability of *P. aeruginosa* to cause infections and significantly decrease the immune response of the host while increasing antibiotic resistance of the pathogen ≥ 1000-fold.^13,14^ Moreover, *P. aeruginosa* may also employ numerous additional virulence factors that allow this infectious prokaryote to adapt to the environment of the host and to evade the immune system. Importantly, upon colonization, *P. aeruginosa* undertakes a transcriptional reprogramming in order to transition to a biofilm– and virulence factor-producing mode. This process is accomplished chiefly by TCSs that function to control the transcription of genes responsible for the formation and dispersal of biofilms as well as several essential virulence factors.^19–22^

One such critical TCS in *P. aeruginosa* is BqsRS, first discovered as a regulator of biofilm dispersal in PAO1,^23^ then later found to sense extracellular Fe^2+^ and to promote resistance to cationic stress in PA14.^24^ BqsRS, also known as CarRS, has also been reported to be induced highly by Ca^2+^, to function as a regulator of Ca^2+^ homoeostasis, and to control Ca^2+^– dependent pyocyanin production and tobramycin resistance.^25^ *Pa*BqsRS is a canonical two-component system composed of a membrane-bound HK, BqsS, and its cognate RR, BqsR. Several lines of evidence have shown that the membrane HK *Pa*BqsS recognizes Fe^2+^ at μM metal concentrations as well as Ca^2+^ at mM concentrations, both of which are physiologically-relevant concentrations of metal cations within the lungs of CF patients.^26–31^ Importantly, *P. aeruginosa* deletion mutants of *bqsS* and/or *bqsR* are defective in the production of short chain quorum sensing signals and rhamnolipids, glycolipid biosurfactants necessary to maintain biofilm architecture.^23^ This BqsRS-mediated alteration in biofilm maintenance ultimately leads to biofilm dispersal that is commonly associated with the spread of infection, indicating that the BqsRS likely plays a critical regulatory role in *P. aeruginosa* pathogenicity.^32^ Furthermore, studies using the motif search program MEME^33^ coupled with electrophoretic mobility shift assays (EMSAs) discovered that the response regulator *Pa*BqsR binds within a tandem repeat DNA sequence 5’-TTAAG(N)_6_-TTAAG-3’ (also known as the BqsR box).^24^ This sequence is upstream of over 100 genes in *P. aeruginosa*, including those involved in metal transport, lipopolysaccharide modulation, polyamine synthesis and transport, and even antibiotic resistance, suggesting that this TCS could be a potential drug target and emphasizing the importance of investigating the structural properties of this system, its regulated genes, and its mechanism of action.^24,34,35^

We have recently characterized intact and truncated *Pa*BqsS to understand its function and its mechanism of metal selectivity.^36^ Specifically, we showed that intact, detergent-solubilized *Pa*BqsS functions as a homodimer and binds a single Fe^2+^ ion within a dimeric interface of the *Pa*BqsS periplasmic domain. X-ray absorption spectroscopy (XAS) and site-directed mutagenesis demonstrated that this Fe^2+^ ion was bound in an octahedral geometry and coordinated by two critical Glu residues, supporting previous transcriptomics data.^24^ We then characterized the ATP hydrolysis of intact *Pa*BqsS and its truncated cytoplasmic domain, and we showed that intact *Pa*BqsS was selectively and exclusively stimulated by the binding of Fe^2+^ in its periplasmic sensor domain. This study was the first to characterize an intact HK bound to its cognate metal. However, many questions remained regarding the structure, the function, and the DNA-binding mechanism of the cognate RR of this system, *Pa*BqsR.

In this study, we have determined the structural and the functional properties of the intact, activated, and truncated forms of the *P. aeruginosa* response regulator BqsR. We have solved the X-ray crystal structure of the N-terminal *Pa*BqsR receiver domain to 1.3 Å resolution, we have determined the structure of the C-terminal *Pa*BqsR DNA-binding domain (DBD) using nuclear magnetic resonance (NMR), and we have modeled the *Pa*BqsR-DNA interactions using AlphaFold. Excitingly, we have found a *Pa*BqsR DNA-binding site upstream of the *feo* operon that encodes for the primary Fe^2+^ transport machinery in *P. aeruginosa*. Using *in vitro* DNA-binding assays, we show that *Pa*BqsR can bind upstream of *feo* and within the BqsR box that overlaps the FUR box that regulates this operon in iron-replete conditions. Correlating with these *in vitro* observations, transcriptomic changes in Δ*bqsR* PAO1 show that *feo* is among the most differentially regulated genes by *Pa*BqsR, making *feo* one of the primary targets of this TCS. Moreover, we demonstrate that *Pa*BqsR regulation is dynamic and responds to Fe^2+^ availability in PAO1, likely through the ability of *Pa*BqsR to bind Fe^2+^ itself, which has not been observed in other OmpR-like response regulators, representing an additional layer of regulation in this system. Taken together, these data elucidate the structure and the DNA-binding properties of *Pa*BqsR for the first time, define its transcriptional regulon, and reveal an important function for BqsRS in controlling *P. aeruginosa* Fe^2+^ acquisition.

## RESULTS

### The crystal structure of the N-terminal domain of *Pa*BqsR reveals an OmpR/PhoB-like phospho-acceptor domain

To determine the structural and biophysical properties of *P. aeruginosa* BqsR (*Pa*BqsR), we cloned, expressed, and purified the intact RR to homogeneity (Fig. S1a). At low concentrations (<< 1 mM), *Pa*BqsR migrated via gel filtration as a monomer, although at high concentrations (*ca.* 1 mM) this oligomerization shifted to a dimeric species (Fig. S1b), suggesting this protein is in dynamic equilibrium between monomer and dimer. Using both species of protein, we set up an extensive series of crystallization screens, optimized the obtained crystals, and tested for diffraction. All crystals indexed to the same space group, and our highest resolution was resolved to 1.3 Å (Table S1). However, a Matthew’s analysis of the lattice parameters was inconsistent with the presence of intact *Pa*BqsR. Guided by sequence conservation amongst other similar RRs, we generated independent AlphaFold models of the N-terminal receiver and C-terminal DNA-binding domains (DBD) of *Pa*BqsR, and we used those as search models for molecular replacement (MR). The N-terminal receiver domain alone produced a definitive MR solution, whereas the C-terminal DBD did not. Subsequent refinement converged to a final model (*R*_w_ = 0.158, *R*_f_ = 0.175) in which only residues 1-123 (chiefly comprising the N-terminal receiver domain) are present (Table S1; PDB ID 8GC6). Despite exhaustive attempts to crystallize the intact, wildtype (WT) *Pa*BqsR protein, we were unable to do so, and all conditions produced crystals of the proteolyzed N-terminal receiver domain alone, likely due to the high dynamics in the linker region connecting this domain to the C-terminal DBD.^37^

The N-terminal *Pa*BqsR receiver domain has strong structural homology to phosphor-acceptor domains of the family of OmpR/PhoB-like RRs (Fig. 2a-c). In particular, our new structure reveals that *Pa*BqsR possesses a canonical (βα)_5_ response regulator assembly that consists of a central five-stranded parallel β-sheet surrounded by five α-helices (Fig. 2a,c). This topology is very common for the OmpR/PhoB-like family, and superpositioning reveals strong structural homology (Cα RMSDs < 1.0 Å) to other receiver domains found in bacterial RRs such as PhoP, ArlR, and KdpE (Fig. S2). One end of the receiver domain is highly negative in its electrostatic surface (likely driving interactions with its partner HK *Pa*BqsS), and the key, conserved phospho-accepting residue Asp^51^ stretches out to the surface of this part of the protein, likely to facilitate reaction with the phosphorylated HK to catalyze phospho-relay (Fig. 2a,c). In contrast, the dimerization of *Pa*BqsR is facilitated through a conserved dimerization domain comprising the α4–β5–α5 region (Fig. 2c) that is of mixed electrostatic polarity and ultimately links to the C-terminal DBD through a flexible linker. Once dimerized, *Pa*BqsR (and most response regulators) are expected to create a positively-charged surface that helps facilitate (in part) the dimeric response regulator to interact with the negatively charged backbone of DNA.^5^ Unfortunately, in WT *Pa*BqsR, the dynamics of the flexible linker make this protein prone to spontaneous proteolysis over time in the crystallization drop, which could not be avoided despite numerous efforts. Thus, a different approach was needed to characterize the C-terminal DBD.

**Figure 2.**
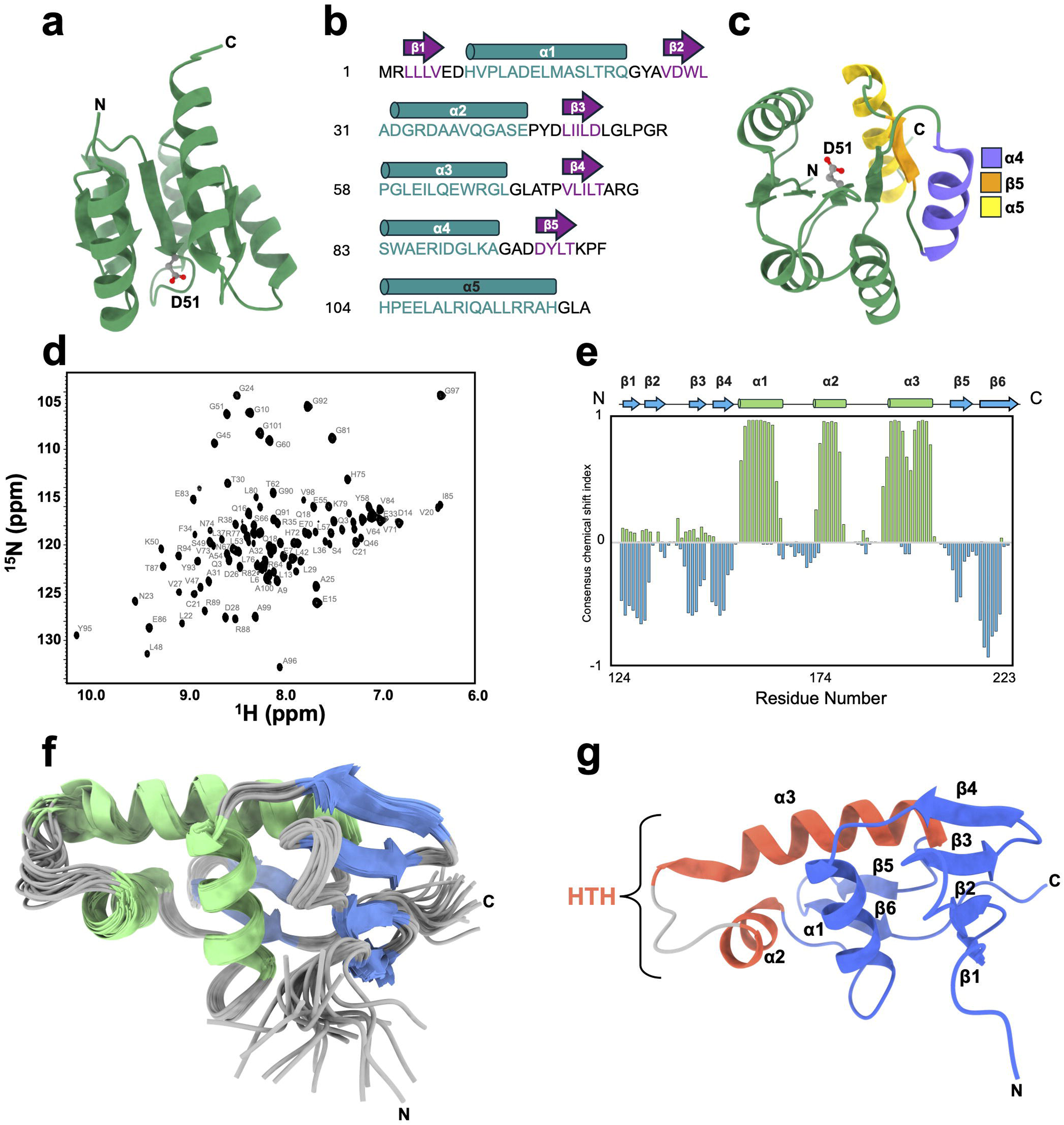
Structural analysis of *Pa*BqsR. **a**. The crystal structure of the *Pa*BqsR N-terminal receiver domain (residues 1-123) reveals an OmpR/PhoB family canonical (βα)_5_ response regulator assembly with a central five-stranded parallel β-sheet surrounded by five α-helices. **b**. Cartoon topology of the *Pa*BqsR N-terminal receiver domain. **c**. Dimer formation for OmpR/PhoB response regulators is generally mediated by the α4–β5–α5 interface, highlighted in purple, orange, and yellow, respectively. **d**. Assigned 2D ^1^H-^15^N HSQC NMR spectrum of *Pa*BqsR DBD. **e**. Consensus chemical shift index (CSI) (Hα, Cα, and Cβ and C0) of PaBqsR DBD, calculated using the program PECAN.^92^ Values nearing +1 indicate α-helical structure, values near 0 indicate unstructures regions/random coil, and values nearing –1 indicate β-strand structure.The cartoon at the top indicates the predicted toplogy of the *Pa*BqsR DBD based on CSI values derived from NMR experiments. **f**. Superpositioning of the 20 lowest-energy refined structures of the *Pa*BqsR DBD (residues 124-223). Labeled are the α-helices (green) and β-strands (blue). **g**. Ribbon diagram of the lowest-energy conformer of the PaBqsR DBD, which contains a helix-turn-helix (HTH) topology common to the OmpR/PhoB-like family. The helix-turn-helix (HTH) motif is colored in salmon and labeled. In all cases, ‘N’ and ‘C’ represent the N– and C-termini in the structure, respectively.

#### The C-terminal DNA-binding domain (DBD) of *Pa*BqsR bears a helix-turn-helix (HTH) topology

As we failed to crystallize the intact structure of *Pa*BqsR, we cloned, expressed, and purified to homogeneity a construct that consisted only of the C-terminal *Pa*BqsR DBD (Fig. S1a). Gel filtration experiments demonstrate that the DBD maintains a monomeric oligomerization at both low and high concentrations (Fig. S1b), in contrast with the WT protein but consistent with the dimerization domain being located in the *Pa*BqsR N-terminal domain. Despite exhaustive attempts, we were unable to produce crystals with this protein. Therefore, we instead turned to nuclear magnetic resonance (NMR) to determine the structure of the DBD. We first collected high-quality 2D ^1^H-^15^N heteronuclear single quantum coherence (HSQC) spectra of the *Pa*BqsR DBD (Fig. 2d) that indicated folded protein based on its well-dispersed amide signals. We then tested several pH ranges to optimize the intensity of the ^1^H and ^15^N chemical shifts (pH 5.5 was found to be optimal), we confirmed that the protein was stable for several weeks at this pH, and we used 2D NMR data collected at this pH for backbone amide assignments. These established NMR conditions ultimately allowed for assignment of all backbone amide signals except Met^1^ (99 % of the non-proline backbone amide signals) (Fig. 2d).

To explore the secondary structure of the *Pa*BqsR DBD, Cα chemical shift indices (CSI) based on the assignment of triple resonance spectra were analyzed (Fig 2e), indicating that the secondary structure of the DBD is composed of a N-terminal set of 4 β-strands (β1, Leu^128^–Ala^131^; β2, Leu^133^–Leu^135^; β3, Cys^141^–Leu^144^; β4, Ala^147^–Asp^150^) followed by three α-helices (α1, Ala^153^–Met^163^; α2, Lys^172^–Leu^179^; α3, Glu^185^–Leu^202^). The DBD then terminates with two β-strands (β5, Ile^207^–Arg^210^; β6, Gly^214^–Tyr^217^) suggesting the presence of a short β-hairpin, a structural feature in DBDs that has been shown to be important in facilitating the interaction between bacterial RRs (such as PhoP, ArlR, and KdpE) and their DNA targets.^38–42^ In order to confirm this architecture, the tertiary structure of the *Pa*BqsR DBD was determined by preparing ^15^N– and ^15^N/^13^C-isotopically enriched protein sample and acquiring 3D ^15^N-edited nuclear Overhauser effect (NOE) spectra.^43,44^ After data acquisition, structural calculations were carried out using a total of 1912 interproton distance restraints derived from NOE data, 48 hydrogen bond restraints, and 148 dihedral restraints based on backbone chemical shifts (Table S2). An ensemble of 20 refined *Pa*BqsR DBD structures was ultimately generated that exhibited good convergence based on root-mean-square deviations (RMSDs) (0.93 Å ± 0.11 Å for backbone heavy atoms) (Fig. 2f, Table S2) (PDB ID 9FYA).

The tertiary structure of the *Pa*BqsR C-terminal DBD is wholly consistent with the CSI analysis and displays a helix-turn-helix (HTH) DNA-binding topology common to the OmpR/PhoB RR family (Fig 2e-g). The N-terminal β-sheet interacts with helix α1 through hydrophobic interactions while π-stacking interactions between the aromatic rings of Phe162 and Tyr215 bring the N-terminal β-sheet in proximity to the C-terminus. The three α-helices (α1-α3) form the trihelical motif of the HTH, with α3 representing the longest and most polar helix that interacts with the DNA major groove (*vide infra*). This overall HTH fold is common to the DBDs of the OmpR/PhoB RR family, and structural superpositioning of the entire DBD reveals strong structural homology (Cα RMSDs < 1.3 Å) to other DBDs found in bacterial RRs such as PhoP, ArlR, and KdpE (Fig. S3).^38–42^ Additionally, the *Pa*BqsR C-terminal DBD ends in a critical β-hairpin that has been shown in other RR DBDs to be important in DNA binding (*vide infra*).^38–42^ However, subtle differences in the DNA-binding helices and this key β-hairpin generally drive specificity and differentiate the gene targets of these RRs, which we then explored.

#### In vitro DNA-binding assays reveal that *Pa*BqsR binds upstream of the *feo* operon in PAO1

As we previously demonstrated that the sensor HK of the BqsRS system, *Pa*BqsS, binds and forms a stable interaction with Fe^2+^,^36^ we hypothesized that one (or several) of the gene targets of this TCS would be involved in iron homeostasis, an essential component for bacterial survival and a critical factor in biofilm formation and virulence of *P. aeruginosa*.^32,45,46^ Prior work defined the BqsR box sequence as 5’-TTAAG(N)_6_-TTAAG-3’^24^, so we first undertook a directed search of a limited portion of the PAO1 genome for this DNA sequence, searching for its presence upstream of genes of interest such as those involved in spermidine synthesis,^47^ Fe^3+^ reduction,^48,49^ and Fe^2+^ acquisition^50^ (Table S3). Given that *Pa*BqsR can sense Fe^2+^ directly, we were particularly interested in whether *Pa*BqsR could target the *feo* operon that encodes for the primary Fe^2+^ transport machinery in PAO1. Interestingly, we observe a putative BqsR box upstream of the PAO1 *feoA* start site, overlapping with a putative ferric uptake regulator (FUR) box (a global, negative iron regulator), but distal from, and independent of, a putative anaerobic transcriptional regulator (ANR) box (a global regulator of anaerobic genes) (Fig. 3a). These results suggested that *Pa*BqsR could regulate *feo* expression in PAO1, which we then tested *in vitro*.

**Figure 3.**
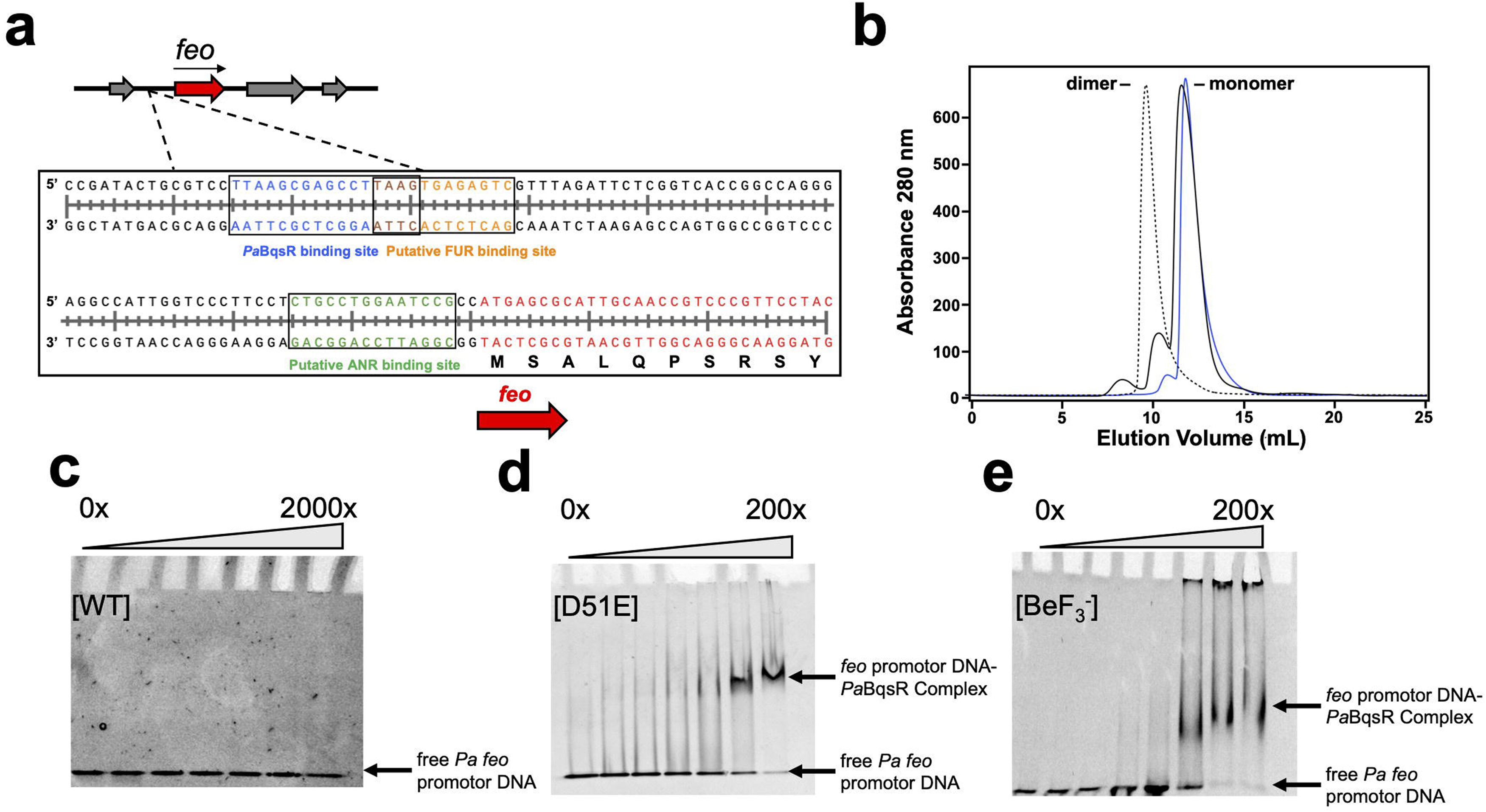
*Pa*BqsR binds upstream of the *feo* operon in PAO1. **a**. A putative BqsR box sequence (5’-TTAAG(N)_6_-TTAAG-3’; blue) is present in PAO1 upstream of *feo*, overlaps with a putative ferric uptake regulator (FUR) box (orange), and is upstream of the anaerobic transcriptional regulator (ANR) box (green), adjacent to the start site of the *feo* operon (red). **b**. Size exclusion chromatography (SEC) profiles of WT *Pa*BqsR (black), D51E *Pa*BqsR (blue), and BeF ^-^-activated *Pa*BqsR (dashed). BeF ^-^-activated PaBqsR is a dimer, while WT and D51E *Pa*BqsR are both monomers at low concentrations (<< 1 mM). **c**. EMSAs with WT *Pa*BqsR and the *feo* duplex. **d**. EMSAs with D51E *Pa*BqsR and the *feo* duplex. **e**. EMSAs with the BeF ^-^-activated *Pa*BqsR and the *feo* duplex. All EMSAs were performed in triplicate with 0.5 µM DNA.

To determine whether *Pa*BqsR could bind to the BqsR box upstream of the *feo* operon, we used *in vitro* electrophoretic mobility shift assays (EMSAs). To do so, we tested binding using a 21 base-pair (bp) DNA duplex corresponding to minimal region needed for protein binding (Table S4). Intact, monomeric, WT *Pa*BqsR (Fig. 3b) failed to bind to this duplex when tested across a range of concentrations from 0-2000 mol. eq. protein:DNA (Fig. 3c). This behavior was expected, as the canonical TCS paradigm indicates that the RR must either be phosphorylated or pseudo-phosphorylated to facilitate DNA binding.^7^ Pseudo-phosphorylation can be achieved by producing an Asp-to-Glu variant or by reaction with BeF ^-^ that binds to the phosphorylatable Asp residue in the RR and mimics the presence of a phosphate group. Canonically, the addition of phosphate to the conserved Asp in the receiver domain is thought to stabilize a distinct conformational and dynamical state that is allosterically communicated to the C-terminal DNA-binding to enhance its DNA-binding affinity.^51–53^ It is reported that in inactive states, the receiver domain and DNA-binding domain interact in such a manner that the DNA-binding domain mobility is constrained, leading to lower DNA affinity.^51,54,55^ The Asp-to-Glu variant in the receiver domain has been shown to approximate the steric presence of a phosphate group attached to Asp, although true binding enhancements often require actual phosphorylation that Glu cannot induce.^56^ Nonetheless, it is often successfully used as a phospho-mimic for response regulators.^55–57^ In contrast to Glu, BeF ^-^ binds to Asp in a similar geometry to phosphate, often inducing a conformational and dynamical state that greater mimics the naturally phosphorylated response regulator.^58–60^ Given that both methods are commonly used, we tested both approaches. We first used mutagenesis to create a *Pa*BqsR D51E variant that we expressed and purified to homogeneity (Figs. 3b, S1b). Interestingly, while this protein remained monomeric in solution (Figs. 3b, S1b), the D51E variant showed clear and distinct binding to the *feo* duplex (Fig. 3d), demonstrating that protein dimerization in the absence of DNA is not a pre-requisite for intact *Pa*BqsR to bind to DNA, and that pseudo-phosphorylation enhances the *Pa*BqsR-DNA interaction. We then generated the pseudo-phosphorylated, WT BeF ^-^-activated form of *Pa*BqsR that was clearly dimeric in solution (Figs. 3b, S1b). The BeF ^-^-activated phospho-mimetic of *Pa*BqsR also bound to the *feo* duplex with an apparent affinity nearly identical to the D51E variant (Fig. 3e). In order to rule out any effects on DNA affinity based on DNA size, we performed EMSAs with a larger DNA duplex (> twice the size) containing the BqsR box, and a similar apparent affinity was observed (Fig. S4e,f). Control EMSAs using bovine serum albumin (BSA) in place of *Pa*BqsR failed to bind to the *feo* duplex, demonstrating this interaction to be *Pa*BqsR-specific (Fig. S4d). Additionally, we tested whether the monomeric *Pa*BqsR C-terminal DBD alone could bind to DNA. Indeed, we observed binding to the *feo* duplex with only the DBD (Fig. S4a), but the apparent binding was 2-3 fold weaker than that of either D51E or the BeF ^-^-activated form of full-length *Pa*BqsR (Fig. 3), indicating that the presence of the N-terminal receiver domain contributes to the binding affinity of this RR to its DNA sequence. Furthermore, EMSAs performed using a scrambled BqsR box elicited no binding (Fig. S4g), demonstrating selectivity for this DNA sequence. Taken together, these results demonstrate that *Pa*BqsR may target the *feo* upstream region *in vitro*, suggesting that this TCS is a regulator of a critical iron uptake system frequently used by *P. aeruginosa* at the host-pathogen interface.^61^

#### Modeling of *Pa*BqsR and the PAO1 *feo* operon illustrates a unique mode of DNA binding

We next sought to use structural approaches to understand how *Pa*BqsR binds to the upstream *feo* region. To do so, we first tried to crystallize both *Pa*BqsR D51E and the BeF ^-^-activated forms of *Pa*BqsR in the presence of the *feo* upstream region duplex modified to include an additional G-C overhang, which has been shown to increase crystallization propensity of other DNA-protein complexes.^62^ While we were able to generate crystals of both complexes, these crystals only diffracted weakly and anisotropically despite extensive optimization. We then turned to AlphaFold to model the *Pa*BqsR-*feo* interaction (Fig. 4a, Fig. S5, S6), as we already possessed experimentally-determined structural information on both domains of *Pa*BqsR. Surprisingly, there is little consensus on how exactly OmpR/PhoB-like RRs recognize and bind to their target DNA sequences. For example, the DBDs of PhoB, PmrA, or RstA bind DNA asymmetrically,^38,40,42^ whereas the DBD of OmpR binds to various DNA sequences in either a symmetric or asymmetric orientation,^39,63,64^ further underscoring the importance of investigating the *Pa*BqsR DNA binding. Consistent with the behavior of the BeF ^-^-activated form of WT *Pa*BqsR, AlphaFold predicts binding of *Pa*BqsR to the *feo* upstream region as a dimer (Fig. 4a), as no binding is predicted when only a single copy of *Pa*BqsR is provided as an input. Intriguingly, modeling predicts that dimerization at the N-terminal phospho-acceptor domain occurs in an asymmetric manner, and this dimerization orientation may be facilitated by the phosphorylation of Asp^51^ (Fig. 4a). Importantly, as predicted based on NMR data (Fig. 2), α3 of the HTH motif of the C-terminal DBD binds within the major groove of DNA (Fig. 4a) and within the variable nucleotide region and upstream (towards the 5’ end of the sense strand) of the conserved 5’-TTAAG-3’ sequence (Fig. 4a). Basic residues of *Pa*BqsR such as Lys^172^, His^194^, His^197^, Arg^199^, Arg^200^, and Arg^204^ create a positively-charged surface that includes the exposed face of α3 on each *Pa*BqsR protomer and extends across the dimer (Fig. 4a-c). This important region contacts the DNA backbone and helps place α3 into the major groove of the DNA (Fig. 4b-d) with the sidechains of Lys^172^, Arg^199^, and Arg^200^ making contact with the DNA phosphates (with the closest sidechain-phosphate distances of 2.72 Å, 2.72 Å, and 2.76 Å, respectively) (Fig. 4d, inset). Intriguingly, the terminating β-hairpin of each dimer contains Arg^211^ that appears to “anchor” *Pa*BqsR within the BqsR box through three specific nucleobases of the tandem repeat: Arg^211^ on the first protomer hydrogen bonds with T^33^ and A^34^ on the sense strand and A^24^ on the antisense strand, while Arg^211^ on the second protomer hydrogen bonds with A^36^ on the sense strand and T^22^ and A^23^ on the antisense strand (Fig. 4d, inset). Thus, modeling suggests that at least 7 different amino acids in the DBD facilitate contacts between each *Pa*BqsR and DNA, with Lys^172^, Arg^199^, Arg^200^, and Arg^211^ initially hypothesized to play the most important roles in recognizing the BqsR box.

**Figure 4.**
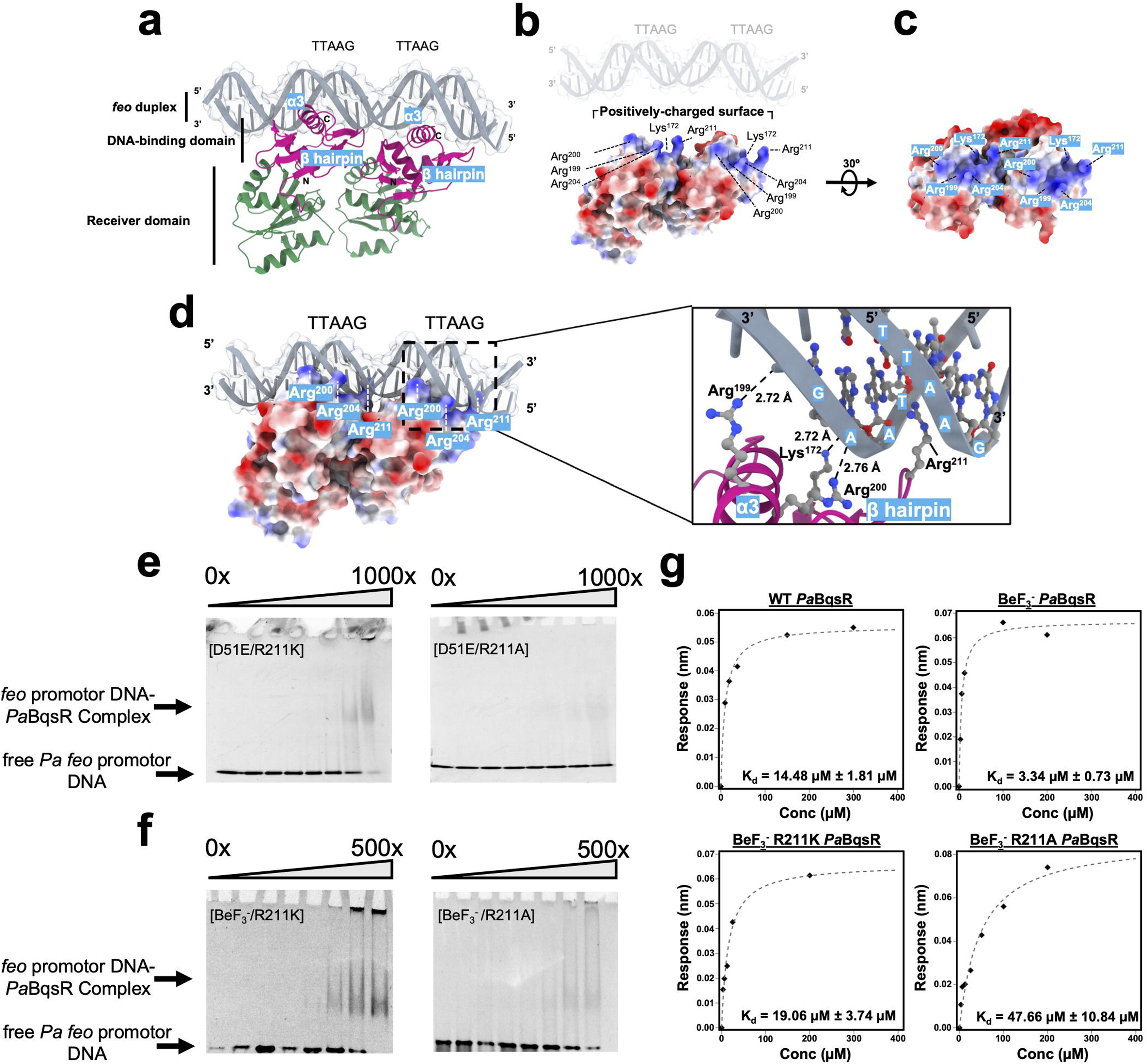
Modeling of *Pa*BqsR binding to the *feo* duplex. **a**. AlphaFold model of WT, intact *Pa*BqsR bound to the *feo* upstream duplex. The region in green is the N-terminal receiver domain, while the region in purple is the C-terminal DNA-binding domain (DBD). **b,c**. An electrostatic surface comprising Lys^172^, His^194^, His^197^, Arg^199^, Arg^200^, and Arg^204^ extends across the dimer and utilizes the exposed α3 of each *Pa*BqsR monomer to bind within the DNA major groove. **d**. Lys^172^, Arg^199^, Arg^200^, and Arg^211^ directly interact with the BqsR box, with residue Arg^211^ forming an anchor that π-stacks into each half of the tandem repeat, providing specificity. In all cases, ‘N’ and ‘C’ represent the N– and C-termini in the structure, respectively. **e**. EMSAs of *Pa*BqsR variants (D51E/R211K, D51E/R211A) in complex with the PAO1 *feo* upstream region indicates that the D51E/R211K variant has a lowered affinity to the PAO1 *feo* upstream region, while the D51E/R211A variant fails to bind to the PAO1 *feo* upstream region at the same ratio. **f**. EMSAs of BeF ^-^-activated *Pa*BqsR variants (R211K, R211A) in complex with the PAO1 *feo* upstream region indicates that both the BeF_3_^-^/R211K variant and the the BeF ^-^/R211A variant have lowered affinity to the PAO1 *feo* upstream region, albeit stronger than the D51E double variants. All EMSAs were performed in triplicate with 0.5 µM DNA. **g**. BLI-based binding curve showing an apparent binding affinity (*K_d_*) of 14.48 ± 1.81 µM for WT *Pa*BqsR and the *feo* upstream region (top left), 3.34 ± 0.73 µM for BeF ^-^-activated *Pa*BqsR and the *feo* upstream region (top right), 19.06 ± 3.74 µM for BeF ^-^-activated R211K *Pa*BqsR and the *feo* upstream region (bottom left), and 47.66 ± 10.84 µM for BeF ^-^-activated R211A *Pa*BqsR and the *feo* upstream region.

To understand whether the intercalating β-hairpin residues are conserved across OmpR-like RRs, and to test the hypothesis that Arg^211^ “anchors” the RR to the *feo* upstream region, we performed bioinformatics, site-directed mutagenesis, and binding studies using both EMSAs and biolayer interferometry (BLI). Several biochemically-characterized RRs conserve the HTH and β-hairpin structural motifs while having poor sequence conservation (Figs. S7). Interestingly, Arg^211^ in the *Pa*BqsR DBD lacks conservation across OmpR-like RRs (Fig. S7), so to test the importance of Arg^211^ in mediating protein-DNA interactions, we expressed, purified, and tested for DNA binding several *Pa*BqsR variants (Fig. S8): R211A (does not dimerize and abrogates the positively-charged locus in this position), R211K (does not dimerize and maintains the positively-charged locus in this position), D51E/R211A (mimics activation but abrogates the positively-charged locus in this position), and D51E/R211K (mimics activation and maintains the positively-charged locus in this position). To confirm that no major change in secondary structure occurred for these variants, we used circular dichroism (CD) spectroscopy that revealed all variant proteins to have a similar secondary structure compared to the WT protein (Fig. S9). As expected, the inactivated R211A and R211K variants failed to bind to the *feo* upstream region. In contrast, for the pseudo-activated D51E/R211K variant, EMSAS showed that binding still occurs, albeit weaker (>5-fold diminished) compared to the D51E variant alone (Fig. 4e), while the pseudo-activated D51E/R211A variant did not bind (Fig. 4e). However, it should be noted that the D51E variants only functions as weak phosphomimics, as these variants remain monomeric in solution, which does not recapitulate the dimerization observed in the native response regulator as well as the BeF_3_^-^-activated protein. To explore the contribution of Arg^211^ to DNA binding in the context of protein dimerization, both the R211K and R211A variants were BeF_3_^-^-activated and analyzed by EMSAs. The BeF_3_^-^-activated R211K variant displayed weaker qualitative binding when compared to BeF ^-^-activated WT *Pa*BqsR, but stronger qualitative binding compared to the D51E/R211K variant as expected (Fig. 4f). Moreover, the BeF ^-^-activated R211A variant displayed an even weaker qualitative binding than the BeF ^-^-activated R211K variant, although some complex formation was observed unlike the D51E/R211A variant (Fig. 4f). Interestingly, when Arg^211^ is present and we performed EMSAs at high salt concentrations (500 mM NaCl) both D51E and BeF ^-^–activated *Pa*BqsR revealed no perturbations in their apparent affinities (Fig. S4h,i); in contrast, when Arg^211^ was absent and we performed EMSAs at high salt concentrations, both the BeF ^-^-activated R211K and R211A variants revealed a qualitative decrease in affinity (Fig. S4j,k). These data demonstrate that both the presence and the composition of the Arg^211^, with both its positive charge and its planar guanidinium side chain, appear to be the main drivers of affinity between *Pa*BqsR and the *feo* promoter region.

To strengthen this argument by using quantitative measures, we then performed BLI to derive equilibrium dissociation constants (*K_d_*) between the *feo* promoter region and either WT *Pa*BqsR or the dimerized BeF ^-^-activated mimics. First, our BLI-derived binding curves revealed that unactivated WT *Pa*BqsR has only modest affinity to the *feo* upstream region with an apparent *K_d_*of 14.48 ± 1.81 µM (Fig. 4g), comparable to the affinity of other unactivated RRs to their target DNAs.^42,65,66^ As a negative control, BLI was performed with the scrambled BqsR box (*vide supra*), and no binding was observed (Fig, S10). As expected, BeF ^-^ activation significantly increased the affinity of *Pa*BqsR for the *feo* upstream with an observed *K_d_* of 3.34 ± 0.73 µM (Fig. 4g), also comparable to the affinity of other pseudo-activated RRs to their target DNAs.^42,65,66^ Consistent with our EMSAs, loss of Arg^211^ (but maintenance of a positive charge at this position) coupled with protein dimerization resulted in a >5-fold weakening of protein-DNA binding as the BeF ^-^-activated R211K *Pa*BqsR variant exhibited an observed *K_d_* value of 19.06 ± 3.74 µM (Fig. 4g). Furthermore, protein-DNA binding was >10-fold weakened for the BeF ^-^-activated R211A *Pa*BqsR variant, as it exhibited an observed *K_d_* value of 47.66 ± 10.84 µM (Fig. 4g), also consistent with our EMSAs. When taken together, these quantitative data provide support for our AlphaFold model (Fig. 4a-d) and our qualitative EMSAs (Fig. 4e,f), revealing that this terminating β-hairpin Arg^211^ “anchors” *Pa*BqsR to the BqsR box and drives both its affinity and specificity.

#### *Pa*BqsR regulates a substantial portion of the PAO1 genome, including the *feo* operon

To determine the *in vivo* regulatory impact of BqsR in *P. aeruginosa*, we conducted an RNA-seq analysis comparing the transcriptomes of wild type (WT) PAO1 and an isogenic Δ*bqsR* mutant (PRJNA874094). For this experiment, total RNA was isolated from both strains grown aerobically to mid-log phase (12 h) in biofilm minimal 8 (BMM8.5; see Methods) with an added sufficient concentration of iron (4 µM FeSO_4_). Differential expression analysis revealed that BqsR regulates approximately 43 % of the PAO1 transcriptome, with 1311 genes significantly upregulated (log_2_-fold change (LFC) in Δ*bqsR* vs WT ≥1, *p* < 0.05) and 1128 genes significantly downregulated (LFC ≤ –1, *p* < 0.05), indicating that the protein functions as both an activator and repressor of gene expression in *P. aeruginosa* (Fig. 5a). The ability of *Pa*BqsR to act as both a positive and negative transcriptional regulator is not unprecedented, as several other RRs have been reported to have a dual mode of regulation.^21,67–69^ However, previous research has implied that *Pa*BqsR acts only as a classical transcriptional activator when phosphorylated.^24^ This vast array of regulation significantly expands on the previously reported regulatory role of the protein in response to Ca^2+^ addition^25^ and Fe^2+^ shock during anaerobic growth,^24,34^ revealing that BqsR directly or indirectly controls the expression of a large number of genes in *P. aeruginosa*. To understand this large response more clearly, we stratified the differentially regulated genes into three categories: low responders in which the expression was altered between |1| ≤ LFC ≤ |2|, including 1527 genes; moderate responders in which the expression was altered between |2| ≤ LFC ≤ |3|, including 616 genes; and 296 top responders in which the expression was altered with a LFC ≥ |3| (Fig. 5a). To identify *Pa*BqsR-regulated processes and to evaluate the biological significance of this regulation, differentially expressed genes (DEGs) were sorted into general functional categories (Fig. 5b). These categories were assigned based on hand-curated gene annotation (see Methods). Interestingly, Fig. 5b shows that every functional category contained genes regulated by *Pa*BqsR. As evaluated based on both the magnitude of expression changes and the number of DEGs, the most significantly affected processes included iron uptake, phosphate metabolism, sulfur metabolism, and various transport systems. Among them, genes encoding iron sequestration (Feo, Pvd, and Fec), bacterial secretion systems (Type III and Type IV), lipopolysaccharide synthesis and modification, vitamin B_12_ synthesis, and urea metabolism were upregulated, whereas genes encoding phosphate and sulfur metabolism and uptake were downregulated in the absence of *bqsR* (Fig. 5c).

**Figure 5.**
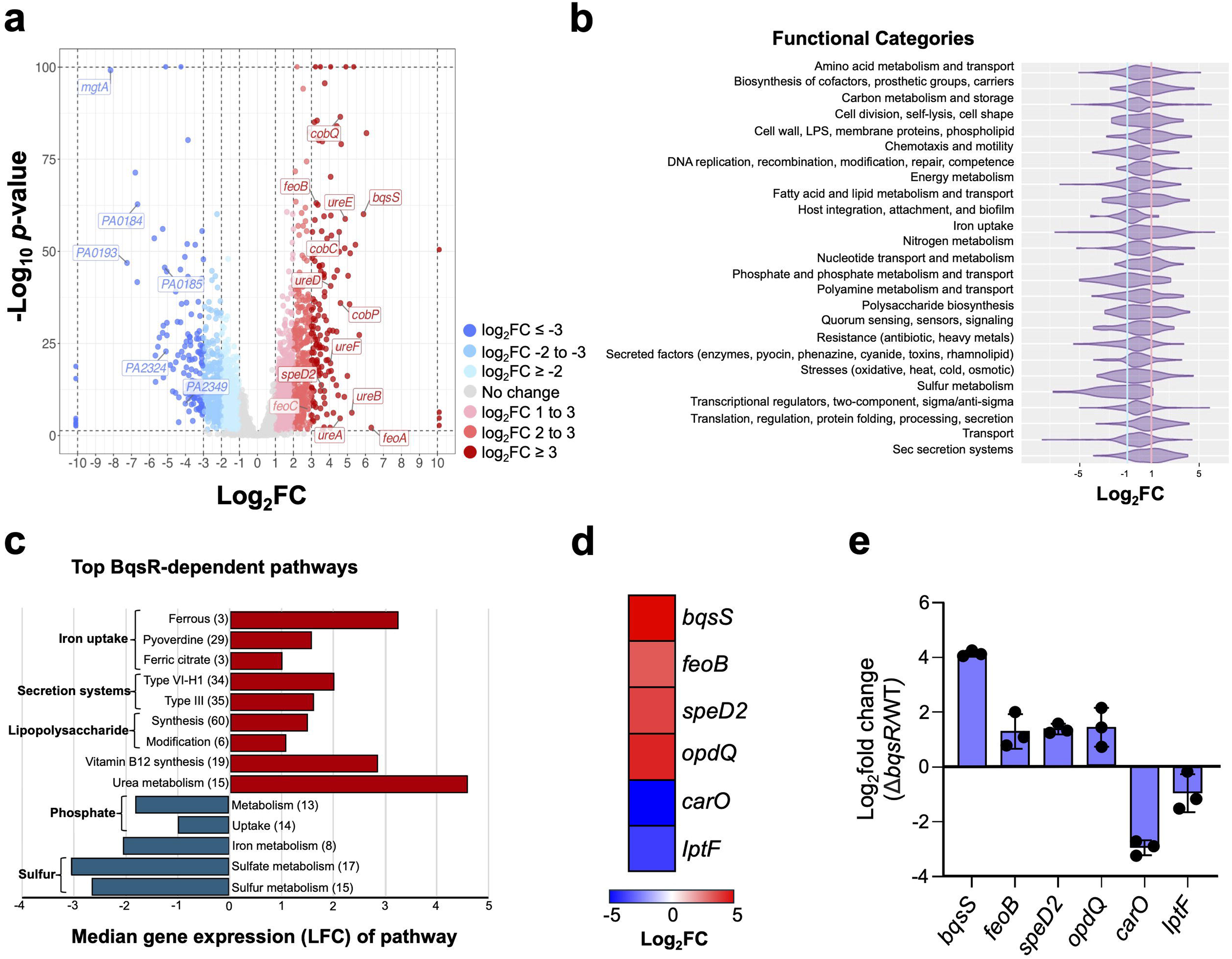
BqsR regulates a significant portion of the PAO1 genome. **a**. Volcano plot depicting the differentially expressed genes (DEGs) in Δ*bqsR* vs PAO1. DEGs (Log_2_FC ≥ |1|, p < 0.05) were stratified based on their log_2_ fold change (LFC) into three categories: low response with the expression changed by LFC |1-2| (light red/blue); moderate response with LFC |2-3| (medium red/blue); and top responders with LFC ≥ |3| (dark red/blue). Genes that showed no change in expression are shown in gray. False discovery rate (p-value) is plotted as –Log_10_ p-value with a cutoff of 1.3 (p = 0.05) indicated by a horizontal dashed line. Genes with LFCs ≥ 10 and/or – log_10_p-values ≥ 100 are depicted along the outermost dashed lines. Genes encoding the most significantly altered pathways are labeled with their gene name or PA number (as annotated in pseudomonas.com). RNAseq-based heat map showing LFC of expression comparisons of *feoABC* and *bqsS* in Δ*bqsR* vs WT PAO1 (inset). **b.** Violin plots showing the functional categories impacted by BqsR in PAO1. The PAO1 functional categories were assigned as described in the Methods. The vertical lines indicate the LFC 1 (red) and –1 (blue). **c**. The top BqsR-dependent pathways within the top differently regulated functional categories. **d**. The top BqsR-dependent DEGs (determined by RNA-seq) that contained a full (5’-TTAAG_NNNNNN_TTAAG-3’) or partial (5’-TTAAG-3’) BqsR box were selected for qPCR validation. **e.** qPCR of the top BqsR-dependent DEGs. The gene transcript abundances in Δ*bqsR* were divided by those in the WT and presented as LFC and plotted as mean (n=3) with standard deviation. Statistical significance was evaluated by using 95 % confidence intervals of LFC shown for each gene and is shown as error bars.

To validate these *Pa*BqsR-dependent transcriptional changes, we then selected six highly *Pa*BqsR-dependent DEGs: *bqsS* (average LFC 5.9), *feoB* (average LFC 3.3)*, speD2* (average LFC 3.7)*, opdQ* (average LFC 4.4) (all negatively regulated by *Pa*BqsR), and *carO* (average LFC –5.0) and *lptF* (average LFC –3.8) (both positively regulated by *Pa*BqsR) (Fig. 5d), and we performed qPCR analysis. The qPCR results (Fig. 5e) confirmed the RNAseq findings that the expression of these genes is regulated by *Pa*BqsR. To evaluate whether the moderate and top responders were direct targets of *Pa*BqsR, we examined the correlation between the magnitude of differential expression and the presence of the BqsR box, previously identified in PA14 (*vide supra*).^24^ For this, we performed a full analysis of the PAO1 genome using the FIMO motif search within the MEME suite.^21^ This analysis identified 103 genes containing the consensus sequence within their putative promoter regions, with only five genes possessing the BqsR box tandem consensus sequence identical to 5’-TAAG(N)_6_-TTAAG-3’. The 103 identified genes showed LFC values ranging from –5.0 to 6.3 and included 50 genes not affected by the deletion under the tested conditions, indicating that direct regulation by *Pa*BqsR may be subtle, that *Pa*BqsR regulation may depend on additional interacting regulators and environmental cues, and that the *Pa*BqsR regulon extends beyond the genes detected in this experiment, further underscoring its role as a global regulator. To understand better the basis of positive and negative regulation by *Pa*BqsR, we selected a subset of genes that were either induced or repressed by *Pa*BqsR, including the six verified by qPCR, and analyzed the relative positions of the BqsR box and the predicted –10/-35 promoter sequences indicated by DeNovoDNA (Fig. S11). The analysis showed that the BqsR box was located downstream of the –10/-35 promoter sequences for the genes positively regulated by *Pa*BqsR (Fig. S11), whereas the genes with negative or both types of *Pa*BqsR regulation showed the BqsR box overlapping with at least one of the –10/-35 elements (Fig. S11). Whether positively or negatively regulated, (such as *carO* or *feo*, respectively (Fig. 5d,e)), EMSAs show that pseudo-phosphorylated *Pa*BqsR reaches binding saturation at similar ratios (Fig. S4 l,m), indicating that direct binding of *Pa*BqsR to the BqsR box can elicit both positive and negative modes of regulation. Although based on a small set of genes, these observations suggest that the position of the BqsR box relative to the predicted promoter provides a basis for two distinct regulatory outcomes of *Pa*BqsR activity and indicates that *Pa*BqsR may act in concert with other transcriptional factors or regulators, such as FUR and ANR, bindings site for both of which are also upstream of the *feo* operon.

One of the most strongly responding pathways regulated by *Pa*BqsR was Fe^2+^ transport encoded by the *feo* operon. While we predicted that BqsR would regulate *feo* based on our *in vitro* results (Fig. 4), we were surprised that our RNA-seq results revealed that *feoA*, *feoB*, and *feoC* were significantly increased in their expression (LFC of 6.3, 3.3, and 2.8, respectively) in the Δ*bqsR* mutant (Fig. 5a), which was validated by qPCR for *feoB* under these conditions (Fig. 5b). This elevated expression of *feoABC* in the absence of BqsR suggests that the protein negatively regulates the *feo* operon in PAO1 grown aerobically. Further, deletion of *bqsR* resulted in a strong upregulation of its cognate sensor, *bqsS* (LFC of 5.9; Fig. 5a,d). This observation is consistent with previous findings in PA14, where BqsR was shown to autoregulate its own expression;^34^ however, in that study, self-regulation of *bqs* was observed to be positive when PA14 was grown under anaerobic conditions in the presence of shocking levels (100 µM) of Fe^2+^. Together, these findings identify a relatively large regulon of *Pa*BqsR and indicate that this TCS has a major role in conditional control of Fe^2+^ acquisition, but the type of regulation initially observed was counter to our expectations.

#### *Pa*BqsR regulation is dynamic and responds to Fe^2+^ availability

While we expected that our RNA-seq data would reveal that the *feo* operon was among the top differentially expressed genes regulated by *Pa*BqsR, we were initially surprised that regulation appeared to be negative at low but sufficient levels of Fe^2+^ (4 µM) (Fig. 5a,d). Since RNA was isolated from aerated aerobic cultures, we hypothesized that Fe^2+^may be oxidized to Fe^3+^ during growth, affecting the output of Fe^2+^-dependent *feo* regulation. To investigate the relationship between Fe^2+^ availability and *feo* expression, we constructed a *feo* promoter (P*^feo^*)-fusion luminescent plasmid-based reporter by cloning a 300-bp DNA fragment upstream of *feoA* immediately prior to the start of the *lux* operon in pMS402, and measured luminescence in tandem with monitoring growth. To maintain Fe reduction for the duration of the experiment, ascorbic acid (AA) was added to a final concentration of 2 mM, which was determined to be the optimal concentration that both maintained maximal Fe^2+^ over time and did not cause a significant growth disadvantage in *P. aeruginosa* (Fig. S12). Additionally, pyruvate (10 mM) was added to the medium to prevent the accumulation of reactive oxygen species generated in the Fenton reaction between ascorbic acid and Fe^2+^ (Fig. S13).^70^ For this set of experiments, the WT and Δ*bqsR* strains harboring the reporter were grown in BMM8.5 (with pyruvate) with no added Fe^2+^ for 8 h and then treated with low (4 µM) or high (100 µM) levels of Fe^2+^ (Fig. 6a). The iron stocks were prepared in the presence of 2 mM ascorbic acid (AA) to minimize oxidation and therefore AA alone served as a control. Prior to Fe^2+^ treatment (0-8 h), the levels of P*^feo^*-driven *lux* expression in Δ*bqsR* were decreased compared to the WT (Fig. 6a), indicating a positive regulation of *feo* by *Pa*BqsR in the absence of iron, likely to obtain Fe^2+^ under iron-depleted conditions. A similar but more pronounced pattern was observed when bacteria were treated with the addition of 4 µM Fe^2+^ in the presence of AA, recapitulating positive regulation of the *feo* operon by *Pa*BqsR (Fig. 6a). However, when PAO1 was treated with 100 µM Fe^2+^ in the presence of AA, the levels of *feo* increased in the mutant compared to the WT, indicating that *feo* is negatively regulated by *Pa*BqsR when PAO1 is exposed to toxic levels of Fe^2+^, likely as a mechanism to mitigate intracellular Fe^2+^ toxicity. Interestingly, 14 h after the addition of 100 µM Fe^2+^ in the presence of AA, *feo* expression becomes positively regulated by *Pa*BqsR (similar to what we observe at 0 and 4 µM Fe^2+^ in the presence of AA), possibly due to the gradual oxidation of Fe^2+^, and/or increased need for Fe^2+^ caused by the repression of *feo* during the first 13 h of growth (Fig. 6a). These results indicate that the regulation of *feo* by *Pa*BqsR is responsive to varying concentrations of Fe^2+^ and that the repression of the Feo system by high levels of Fe^2+^ is likely a mechanism to prevent its toxicity.

**Figure 6.**
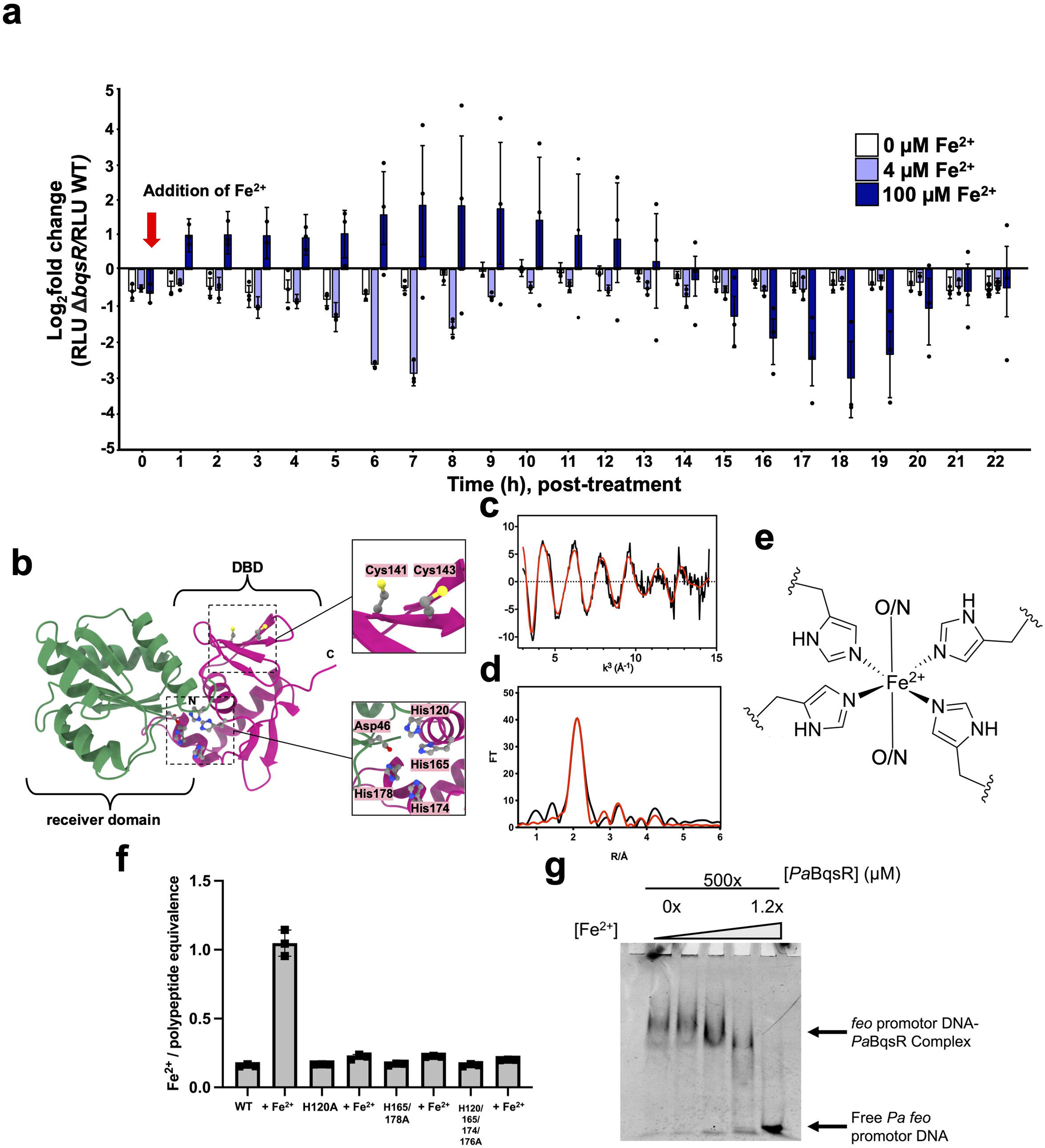
BqsR differentially regulates *feo* expression in response to different Fe^2+^ concentrations. **a.** The WT and Δ*bqsR* strains harboring the *feo* promoter constructs were grown in the absence of iron for 8 h, after which they were treated with Fe^2+^ prepared in ascorbic acid (AA) to achieve final concentrations of 4 µM or 100 µM Fe^2+^ with 2 mM AA. The treatment point is indicated by the red arrow. Cultures treated with AA alone served as the negative control (0 µM Fe^2+^). The relative luminescence units (RLU) were calculated by normalizing luminescence intensity by OD_600_ that were measured hourly. The effect of *bqsR* mutation was assessed by calculating the log_2_fold change (LFC) between average RLU of Δ*bqsR* and WT at each time point. The experiment was performed with three biological replicates per condition. Statistical significance was evaluated by using 95 % confidence intervals, shown as error bars. **b**. AlphaFold model of intact, WT *Pa*BqsR. Highlighted are the His-rich region at the interface of the two domains (*bottom right*) and the Cys-rich region (CxC motif; *top right*) in the DBD. **c**,**d**. EXAFS of *Pa*BqsR bound to Fe^2+^ under anoxic conditions. The processed EXAFS (**c**) and resulting Fourier transformed data (**d**) are consistent with the presence of Fe^2+^ in a His-rich, octahedral geometry. The black traces represent experimental data, while red traces represent EXCURVE-fitted data. **e**. Cartoon representation of the proposed Fe^2+^ coordination of *Pa*BqsR. ‘N’ and ‘C’ represent the N– and C-termini, respectively. **f**. Loss of *Pa*BqsR His residues in panel **c** abrograte Fe^2+^ binding. Error bars indicate ± one standard deviation of the mean from three independent biological replicates. **g**. An increasing presence of Fe^2+^ dissociates the BqsR-*feo* complex. All EMSAs were performed in triplicate with 0.5 µM DNA.

Motivated by these findings, we then tested whether *Pa*BqsR might bind Fe^2+^ directly, which could explain our *in vivo* findings that *Pa*BqsR regulation changes based on Fe^2+^ bioavailability. In our structural characterization of the various domains of *Pa*BqsR (*vide supra*), we noted two regions of the protein conspicuously rich in potential iron-binding residues (a CxC motif in the DBD and a His-rich motif at the domain-domain interface; Fig. 6b), so we tested whether intact, WT *Pa*BqsR could bind Fe^2+^ like that of its HK counterpart, *Pa*BqsS.^36^ Intriguingly, we observed stable binding of a single Fe^2+^ ion to WT *Pa*BqsR after anoxic loading and copious buffer exchanges, as confirmed by ferrozine assays and ICP-MS (Fig. S14). To characterize the identity and local structure of the Fe^2+^ ion bound to WT *Pa*BqsR, we then used X-ray absorption spectroscopy (XAS). The energy of the inflection of the X-ray absorption near-edge structure (XANES) spectrum of Fe^2+^-bound WT *Pa*BqsR as well as its lack of pre-edge features are consistent with an octahedrally-coordinated Fe^2+^ ion (Fig. S15).^71^ Simulations of the extended X-ray absorption fine structure (EXAFS) data of *Pa*BqsR (Fig. 6c,d) reveal only N/O-based environments surrounding the Fe^2+^ ion (Fig. 6e), with *ca.* 4 ligands best fitted to His residues at 2.09 ± 0.02 Å from the Fe^2+^ ion, while *ca.* 2 ligands (perhaps water) are coordinated more weakly and are observed at 2.17 ± 0.02 Å from the Fe^2+^ ion (Table S5). These data definitively rule out coordination of Fe^2+^ at the CxC motif in the C-terminal binding domain and strongly suggest that the His-rich region at the interface of the two domains in *Pa*BqsR (comprising His^119^, His^164^, His^174^, His^178^) binds the Fe^2+^ ion (Fig. 6b). Consistent with these notions, anoxic Fe^2+^ loading with just the C-terminal DBD of *Pa*BqsR failed to show any metal binding. To confirm this His-rich region on *Pa*BqsR is the location of Fe^2+^ binding, we generated, expressed, purified, and tested for Fe^2+^ binding several *Pa*BqsR variants (Fig. S16): H120A, H165/178A, and H120/165/174/178A. CD spectroscopy confirmed a similar protein fold in all variants compared to WT (Fig. S9), and loss of these His residues abrogated Fe^2+^ binding (Fig. 4f), in support of our hypothesis. Importantly, to probe the effects of Fe^2+^ binding on *Pa*BqsR function *in vitro*, EMSAs were performed anoxically with increasing concentrations of Fe^2+^, revealing that the presence of Fe^2+^ facilitates the dissociation of the BqsR-*feo* complex (Fig. 6g). When our *in vivo* observations are taken together with the unprecedented finding that *Pa*BqsR binds Fe^2+^ directly and probits binding to the *feo* upstream region, we believe these data point to a dual mechanism for *Pa*BqsR to regulate Fe^2+^ uptake in PAO1: first via the BqsS sensing axis directly,^36^ and second via the bioavailable pool of Fe^2+^, independent of BqsS. This proposed mechanism would allow PAO1 to respond rapidly to changes in Fe^2+^ levels so as to prevent toxic levels of Fe^2+^ from accumulating within the bacterium, and our observations represent the first example in which a cytosolic RR of a TCS binds and responds to the same signal/stimulus that is sensed extracytoplasmically by its cognate HK.

## DISCUSSION

TCSs are known to play a key role in regulating the production of virulence factors by infectious bacteria,^5,72^ allowing for these pathogens to adapt and to establish infection within diverse environments of host organisms. As just a couple of examples, the *E. coli* PhoPQ TCS alters the ability of *E. coli* to survive when challenged to antibiotics,^73,74^ while the *F. tularensis* QseBC TCS alters bacterial motility for downstream infection.^75^ Germane to this work, the *P. aeruginosa* BqsRS TCS regulates the production of the virulence factor pyocyanin,^25^ the expression of genes involved in biofilm formation,^23^ and the expression of genes involved in antibiotic resistance.^24^ Despite this critical link between TCS function and the establishment of infection,^23^ our understanding of how a membrane HK selects for its cognate stimulus, and how its cognate RR is selective for target genes both remain understudied, particularly for the Fe^2+^-sensing *Pa*BqsRS system. While *Pa*BqsS was recently characterized at the protein level,^36^ its cognate partner, *Pa*BqsR, had not been. Based on sequence homology, *Pa*BqsR was predicted to be an OmpR/PhoB-like RR,^24^ the most common family of RR present in Gram-negative bacteria.^76^ Consistent with this notion, our X-ray crystal structure of the *Pa*BqsR receiver domain reveals a (βα)_5_ response regulator assembly consisting of a central five-stranded parallel β-sheet surrounded by five α-helices with an overall strong structural homology to members of the OmpR-like family. Stretching to the surface of the domain is the Asp residue (Asp^51^) conserved across the OmpR-like family that receives the phosphate moiety from the HK to set into motion the downstream DNA-binding process, mediated by the *Pa*BqsR DBD. Our NMR structure reveals that the *Pa*BqsR DBD comprises a trihelical helix-turn-helix (HTH) fold that is common for a number of DNA-binding proteins.^38–42^ As the DBDs of RRs determine promoter sequence recognition within an organism,^63^ and as our previous work demonstrated that the membrane HK of the BqsRS system, *Pa*BqsS, was selectively activated by Fe^2+^,^36^ we then considered whether *Pa*BqsR might target one or several genes related to Fe^2+^ homeostasis. Indeed, we have now shown that the BqsR box is found in the promoter region of *feo*, and *in vitro* EMSAs combined with BLI data using recombinantly-produced, pseudo-activated *Pa*BqsR confirmed its ability to bind to the upstream region of *feo*, which is dependent on a critical β-hairpin residue in the *Pa*BqsR DBD (Arg^211^) for recognition of the BqsR box, helping define the structural basis of DNA binding by this OmpR-like RR that we then explored *in vivo*.

Consistent with our *in vitro* EMSA results showing *Pa*BqsR binding to the BqsR box upstream of the *feo* operon, we then demonstrated that the *feo* operon is one of the top *Pa*BqsR-regulated pathways *in vivo*. RNA-seq analysis supported by qPCR revealed that, during aerated aerobic growth of PAO1 in the presence of sufficient iron, transcription of the *feo* operon was negatively regulated by *Pa*BqsR. Although *feo* genes have not been reported as a part of the anaerobic BqsSR regulon in response to ≥100 µM Fe^2+^,^24,34^ negative regulation of *feo* by *Pa*BqsR can be concluded from the follow-up study,^77^ where 50 µM Fe^2+^ led to increased *bqsR* expression but decreased *feo* expression, suggesting iron toxicity at these concentrations. To gain further insight into this regulation, we then performed P*^feo^*-*lux* reporter assays in the presence of AA and pyruvate to preserve Fe^2+^ from oxidation and to mitigate oxidative stress, respectively. Whereas the addition of 4 µM Fe^2+^ in the presence of AA showed a decrease in the P*^feo^*-lux expression in the absence of *Pa*BqsR compared to WT, the opposite was observed initially in response to 100 µM Fe^2+^ in the presence of AA; however, after extended growth, this behavior reverted back to the positive pattern regulation of *feo* seen when 4 µM Fe^2+^ was added to the growth media in the presence of AA. These results suggest that *Pa*BqsR switches its regulation of *feo* from positive to negative when Fe^2+^ is present at potentially toxic, high levels. The latter likely occurs in coordination with the global regulator FUR,^78^ the mechanistic details of which will be the focus of future studies. These results are further corroborated by our *in vitro* EMSAs, which indicate that the presence of Fe^2+^ dissociates the BqsR-*feo* complex. These exciting findings suggest that *Pa*BqsR operates in two modes to control *feo* regulation, positive and negative, with a switch occurring at higher Fe^2+^ concentrations. We proposed that this switch is driven by the ability of *Pa*BqsR to bind Fe^2+^ directly, ultimately causing dissociation *of Pa*BqsR from the BqsR box upstream of *feo*, shown here for the first time and unprecedented among OmpR-like RRs to our knowledge. It is noteworthy that *Pa*BqsR-dependent regulation was detected under lower than 10 µM Fe^2+^, which was previously reported to be required for *Pa*BqsR anaerobic expression.^34^ This result may reflect differences in growth conditions (aerobic *versus* anaerobic) or strain background (PAO1 *versus* PA14). Furthermore, elevated *feo* transcript levels have been detected in CF sputum compared to *in vitro* conditions,^77,79,80^ suggesting the importance of Fe^2+^ homeostasis for *P. aeruginosa* survival during infection. This assertion is substantiated by the evidence that Fe^2+^ abundance increases with the progression of CF lung infections, becoming the dominant iron source,^77^ thereby necessitating tight regulation of the Feo system to balance iron acquisition without causing toxicity. Our study is the first to detail this *Pa*BqsR-dependent mechanism controlling the regulation of *feo* and its biological significance, highlighting *Pa*BqsR as a key component in the intricate regulatory network governing iron homeostasis in *P. aeruginosa* (Fig. 7).

**Figure 7.**
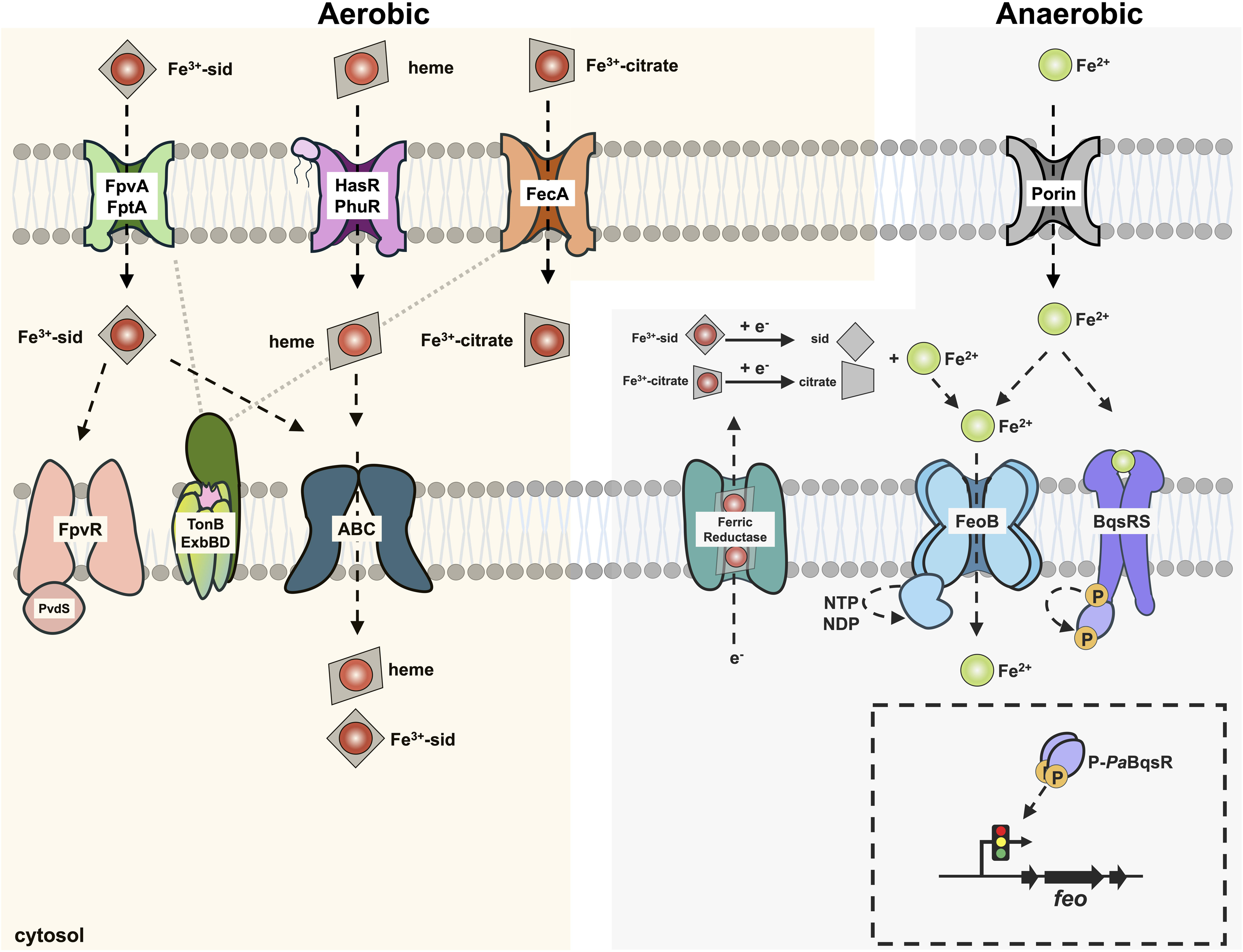
Cartoon of the primary iron homeostasis pathways in *P. aeruginosa* and their relationships to BqsRS. Under oxic conditions, chelated Fe^3+^-siderophore complexes such as pyoverdine are shuttled into the periplasm by outer membrane proteins such as FpvA and FptA. Fe^3+^-citrate uptake is accomplished by FecA. Iron protoporphyrin IX (heme *b*) is sensed and imported by the HasR/PhuR system. These chelated iron complexes are typically brought into the cytosol via ABC transporters, although other mechanism may strip the chelates of the iron itself. Under anoxic conditions, Fe^2+^ is thought to diffuse into the periplasm through outer membrane porins or may be generated in the periplasmic space by ferric reductases. The Feo system then transports Fe^2+^ into the cytosol. The TCS BqsRS directly senses Fe^2+^ and dynamically regulates *feo* (among other genes), which is shown here for the first time.

While previous studies have shown that *Pa*BqsR regulates gene expression during aerobic^25^ and anaerobic^24,34^ growth of *P. aeruginosa* strains, the scope of the regulatory impact of *Pa*BqsR had not been defined. In this work, we identified a large regulon comprising *ca*. 43 % of the PAO1 genome (≥ 2 LFC), 37 % of which belongs to moderate and top responders (≥ 3 LFC), expanding the repertoire of *Pa*BqsR-regulated genes to include iron uptake, urea metabolism, vitamin B_12_ synthesis, lipopolysaccharide (LPS) synthesis/modification, as well as phosphate and sulfur uptake. The previously reported *Pa*BqsR anaerobic regulon in response to Fe^2+^ shock in PA14 was relatively small, including only 46 induced genes.^24,34^ Of these, nine genes (including *carO* and *carP*, which play a role in oxidative stress tolerance and pyocyanin production)^25,81^ were induced and 14 were repressed by *Pa*BqsR during aerated growth with low but sufficient added Fe^2+^. Interestingly, in response to Fe^2+^ shock, *Pa*BqsR induced the expression of the *pmrAB* TCS that controls lipopolysaccharide modification genes but was thought to be only Fe^3+^-responsive.^82^ In contrast, here we show that *Pa*BqsR represses *pmrAB* and lipopolysaccharide modification genes during aerated growth with low but sufficient added Fe^2+^. These findings further underscore that *Pa*BqsR regulation switches from activation to repression depending on both growth conditions and iron concentrations, indicating the capacity of *Pa*BqsR to function as a dynamic global regulator. Considering the physiological significance of the pathways regulated by *Pa*BqsR, this regulatory flexibility likely enhances the ability of *P. aeruginosa* to adapt and to survive within diverse environments and host-associated niches in which nutrients, such as Fe^2+^, may be withheld and dramatically change in their availability. Given this importance, the BqsRS system may represent a very attractive future therapeutic target to treat *P. aeruginosa* infections.

## METHODS

### Materials

All codon-optimized genes used in this study were synthesized and verified by GenScript. Materials used for buffer preparation, protein expression, and protein purification were purchased from standard commercial suppliers and were used as received. Isotopically-enriched ammonium chloride (^15^NH_4_Cl) and glucose (globally ^13^C_6_-labeled) were purchased from Cambridge Isotope Laboratories and used as received. All DNA primers and duplexes used for EMSAs were synthesized and verified by Integrated DNA Technologies. D_2_O was purchased from MilliporeSigma and used as received.

### Bacterial strains and media

Strains and plasmids used in this study are listed in Table S6. *P. aeruginosa* strain PAO1 is the most commonly used non-mucoid strain originally isolated from a wound infection that was sequenced first^83^ and become a reference strain for *Pseudomonas* genetics and functional studies.^84^ All the strains in this study were maintained in 10 % skim milk at −80 °C. For each experiment, bacteria were inoculated from frozen stocks onto LB agar containing the appropriate antibiotic when applicable and grown overnight at 37 °C, from which isolated colonies were used to inoculate precultures. PAO1 and its derivatives were grown in the biofilm minimal medium (BMM)^85^ that was modified and designated BMM8.5 with the following composition: 9.0 mM sodium glutamate, 5 mM glycerol, 0.08 mM MgSO_4_, 0.15 mM NaH_2_PO_4_, 0.34 mM K_2_HPO_4_, 145 mM NaCl, 200 µL trace metals, and 1 mL vitamin solution (per L of medium). The trace-metals solution was prepared with 5.0 g CuSO_4_·5H_2_O, 5.0 g ZnSO_4_·7H_2_O, and 2.0 g MnCl_2_·4H_2_O per liter of 0.83 M hydrochloric acid. The vitamin solution was prepared by dissolving 0.5 g thiamine and 1 mg biotin (per L of the final medium). When relevant, 10 mM sodium pyruvate was supplemented into the BMM8.5 medium. Filter sterilized trace metals, MgSO_4_, and the vitamin solution were added aseptically to autoclaved media and adjusted to a pH of 7.0. This prepared BMM8.5 contains no iron and was designated no-iron-BMM8.5. To prepare BMM8.5 with added iron, FeSO_4_ was prepared as 100x stocks containing either 360 µM or 10 mM iron. To minimize the oxidation of Fe^2+^, 100 mM ascorbic acid was dissolved in the water prior to mixing in FeSO_4_ (1 mM final). The stocks were then filter sterilized and added to the no-iron-BMM8.5 to a final concentration of 4 µM or 100 µM of FeSO_4._

### Protein expression and purification

Codon-optimized DNA blocks encoding for *Pseudomonas aeruginosa* strain PAO1 BqsR (Uniprot identifier Q9I0I1) with either intact (amino acids 1-223, WT *Pa*BqsR), variants H120A, H165/178A, H120/165/174/178A or the C-terminal DNA-binding domain (amino acids 123-223, *Pa*BqsR DBD) modified to encode for an additional C-terminal tobacco etch virus (TEV) protease site (ENLYFQS) were commercially synthesized by Genscript. These DNA sequences were then subcloned into the pET-21a(+) expression vector such that the resulting expression construct was read in-frame with a terminating C-terminal (His)_6_ tag. Generation of the intact D51E, R211A, or R211K variants of *Pa*BqsR was accomplished using site-directed mutagenesis. Mutations to the WT *Pa*BqsR expression plasmid was accomplished by the QuikChange Lightning Multi Site-Directed Mutagenesis kit (Agilent). The following primers were used to introduce the mutated bases (underlined): (D51E forward: 5’-CCCGGCAGACCCAGTTCCAGAATGATCAG-3’; D51E reverse: 5’-CTGATCATTCTGGAACTGGGTCTGCCGGG-3’; R211A forward: 5’-CGAAACCCGTGCTGGTCAAGGTTAC-3’; R211A reverse: 5’-GTAACCTTGACCAGCACGGGTTTCG-3’; R211K forward: 5’-CGAAACCCGTAAAGGTCAAGGTTAC-3’; R211K reverse: 5’-CGAAACCCGT<u>AAA</u>GGTCAAGGTTAC-3’; R211K reverse: 5’-GTAACCTTGACC<u>TTT</u>ACGGGTTTCG-3’). The presence of the appropriate mutation was verified by nanopore DNA sequencing. Each expression plasmid was electroporated into electrocompetent *Escherichia coli* BL21 (DE3) cells, spread onto Luria Broth (LB) agar plates supplemented with 0.1 mg/mL ampicillin (final), and grown overnight at 37 °C. Large-scale expression of all natural-abundance *Pa*BqsR constructs (intact WT, D51E, R211A, R211K, D51E/R211A, D51E/211K, or the DBD) in rich media was accomplished in 6 baffled flasks each containing 1 L sterile LB supplemented with 0.1 mg/mL ampicillin (final) and inoculated with a pre-culture derived from a single bacterial colony. Cells were shaken at 200 RPM until the optical density (OD) at 600 nm (OD_600_) reached *ca*. 0.6-0.8. Cell cultures were then cold-shocked for approximately 2 hours at 4 °C before being induced with 1 mM (final) isopropyl β-D-1-thiogalactopyranoside (IPTG) and incubated at 18 °C overnight with continued shaking. Cells were then harvested by centrifugation for 12 min at 5000×*g* and 4 °C and resuspended with a cellular resuspension buffer (50 mM Tris, pH 7.5, 300 mM NaCl, 10 % (v/v) glycerol, 1 mM TCEP). Immediately prior to cell lysis, solid phenylmethylsulfonyl fluoride (PMSF, *ca.* 50-100 mg) was added to the cellular suspension. Cells were then kept cold in an ice bath while being disrupted via sonication (80 % maximal amplitude, 30 s on pulse, 30 s off pulse, 12 min total time on). The resulting lysate was clarified by ultracentrifugation at 160,000×*g*, 4 °C for 1 hr. The clarified lysate was then purified via immobilized metal affinity chromatography (IMAC) and size-exclusion chromatography (SEC). Briefly, the lysate was injected onto a prepacked 5 mL HisTrap HP column (Cytiva). The column was then washed with 5 column volumes (CVs) of wash buffer (50 mM Tris, pH 7.5, 300 mM NaCl, 10 % (v/v) glycerol, 1 mM TCEP) and eluted stepwise with 7.0 %, 16.7 %, 50.0 %, and 100.0 % (all v/v) respective concentrations of elution buffer (the same composition of the wash buffer plus 300 mM imidazole). Fractions containing *Pa*BqsR were pooled and concentrated using a 10 kDa molecular weight cutoff (MWCO) Amicon spin concentrator. The protein was then buffer exchanged into a TEV cleavage buffer (50 mM Tris, pH 8.0, 200 mM NaCl, 5 % (v/v) glycerol, 1 mM TCEP, 0.5 mM EDTA) by repeated concentration and dilution in the same spin concentrator. A mixture was then made comprising *ca*. 20 μg TEV protease per *ca*. 1 mg protein and allowed to rock at 4 °C overnight. Following tag cleavage, the protein sample was applied to a HiLoad 10/300 GL Superdex 75 preparative SEC column (Cytiva) pre-equilibrated with SEC buffer (50 mM Tris, pH 7.5, 100 mM NaCl, 5 % (v/v) glycerol, 1 mM TCEP) and eluted isocratically. Fractions containing monomeric protein were collected, quantified, and analyzed using 15 % SDS-PAGE and Western blotting. The phosphomimetic *Pa*BqsR-BeF ^−^ complex was generated by incubating WT *Pa*BqsR with 7 mM MgCl_2_, 5.3 mM BeSO_4_, and 35 mM NaF (all final concentrations) prior to further experimentation.

### Crystallization and data collection

Initial crystals of *Pa*BqsR were obtained using sparse-matrix screening applying the sitting-drop vapor-diffusion method. Crystals were subsequently optimized by sitting-drop vapor-diffusion using MiTeGen-XtalQuest Plates with a 1:1 (v:v) of *Pa*BqsR (*ca.* 10 mg/mL) and reservoir solution mixture at room temperature. The precipitant solution consisted of 1.0 M sodium/potassium phosphate at pH 8.2. Medium, three-dimensional, colorless octahedral crystals appeared within 72 hr and reached their maximal sized within 7 days (on average). Crystals were transferred into cryoprotectant, looped, flash-frozen, and stored in liquid N_2_. Data sets were collected on beamlines LS-CAT 21-ID-F at the Advanced Photon Source, Argonne National Laboratory, using a Marmosaic 300 mm CCD detector. Data were processed automatically with Xia2.^86^ Phases were determined via molecular replacement using Phaser in Phenix^87^ with an input of a truncated *Pa*BqsR AlphaFold model (AF-Q9I0I1-F1-v4) (pLDDT score and Predicted aligned error (PAE) for this structure is provided in Fig. S5). Extended model building and refinement cycles were performed in Coot^88^ and Phenix Refine^87^ respectively, and final validations were performed using Phenix Validate.^87^ This final structure, when compared to the initial AlphaFold model, had a Cα RMSD between 117 pruned atom pairs of 0.769 Å. The final model consists of the N-terminal receiver domain of *Pa*BqsR (amino acids 1-123) and has been deposited in the Protein Data Bank (PDB ID 8GC6).

### Nuclear magnetic resonance (NMR) spectroscopy

In order to isotopically-label *Pa*BqsR (intact WT, D51E, or the DBD), the cellular growth and protein production processes were both slightly modified. Briefly, *E. coli* BL21 (DE3) cells containing each expression plasmid grown in minimal media (1x M9 minimal salts, 1 mM MgSO_4_, 1x trace elements, 0.1 mM ZnCl_2_, 0.1 mM CaCl_2_, and 1x MEM vitamin solution) supplemented with ^15^NH_4_Cl, ^13^C_6_-glucose, or both, for global nitrogen and carbon labeling, respectively. Cells were then grown with shaking, induced, harvested, resuspended, and lysed in the same manner as cells grown in rich media (*vide supra*). Isotopically-labeled *Pa*BqsR proteins were purified in the same way as their natural-abundance isotopologues (*vide supra*). Once purified, isotopically-labeled *Pa*BqsR proteins (intact WT, D51E, or the DBD) were then buffer exchanged into an NMR-compatible buffer (50 mM sodium phosphate, pH 5.5, 5 mM NaCl, 1 mM DTT, and 10 % (v/v) D_2_O). All NMR spectra were acquired at 25 °C on either a Bruker 800 MHz spectrometer or a Bruker 600 MHz spectrometer equipped with a cryogenic probe. Protein backbone resonance assignments of the *Pa*BqsR DBD were achieved by standard triple resonance experiments, including HNCA, HBHA(CO)NH, HNCACB, CBCA(CO)NH, and HNCO.^89–93^ Consensus chemical shift index (CSI) (Hα, Cα, and Cβ and C0) of the *Pa*BqsR DBD were calculated using the program PECAN.^94^ Both ^13^C– and ^15^N-labeled three-dimensional nuclear Overhauser effect spectroscopy (NOESY) data were collected in order to calculate the lowest-energy tertiary fold conformers of the *Pa*BqsR DBD. The solution structure of the *Pa*BqsR DBD was calculated using CYANA 3.98.15.^95^ Distance restraints were generated by picking NOE peaks using NMR Analyst^96^ followed by automated NOE cross-peak assignment with CYANA 3.98.15.^95,97^ Upper interproton distance limits of 2.7 Å, 3.3 Å, and 5.0 Å were used for NOE cross-peaks of strong, medium, and weak intensities, respectively. The TALOS+ Server was used to determine backbone dihedral restraints that were incorporated into structural calculations based on amide proton, amide nitrogen, Hα, Cα, Cβ, and carbonyl carbon chemical shifts.^98^ Hydrogen bond restraints were determined based on structure assignments given by the Cα chemical shift indices and strong NOE patterns.^99^ A list of restraint statistics is provided in Table S2. UCSF ChimeraX was employed to prepare structural figures. Out of the 160 calculated conformers, the atomic coordinates of the 20 *Pa*BqsR DBD conformers with the lowest target functions were deposited to the PDB (PDB ID 9FYA). NMR chemical shifts and corresponding structure refinement parameters were deposited in the BMRB database (BMRB accession number 31268).

### Multiple sequence alignments

Response regulators in which the structures of their DNA-binding domains have been experimentally determined were extracted from the PDB, and their sequences were obtained from the Uniprot database. A multiple sequence alignment was constructed through the EMBL MUSCLE program using default parameters.^101^ The resultant alignment was visualized on the MEGAX software.^102^

### Electrophoretic mobility shift assays

Electrophoretic mobility shift assays (EMSAs) were performed in order to test the DNA binding capabilities of *Pa*BqsR to the upstream region of the PAO1 *feo* operon. Increasing concentrations of either intact WT, D51E, R211A, R211K, D51E/R211A, D51E/R211K, or BeF_3_^-^-activated *Pa*BqsR were incubated for 30 minutes at room temperature in SEC buffer with a constant concentration (0.5 µM) of double stranded *feo* DNA containing the BqsR box (Table S4). Analyses were then performed at 4 °C using a 5 % acrylamide gel (0.5 % TB, 29:1 acrylamide:bis-acrylamide). DNA was stained using ethidium bromide and visualized with an ultraviolet (UV) transilluminator.

### Biolayer interferometry

Biolayer interferometry (BLI) experiments were performed using an Octet R2 system (Sartorius) equipped with Octet Anti-HIS (HIS2) biosensors (Sartorius). Briefly, purified WT and variant *Pa*BqsR (15 ng/µL) were immobilized on the HIS2 biosensor surface after hydrating the biosensor for 10 min in SEC buffer. These protein-immobilized biosensors were then submerged into varying concentrations (0–300 µM) of DNA and a reference sample (SEC buffer) for background correction. The BLI responses as a function of time data were recorded and processed using the BLI system-integrated software. *K_d_* values were determined by plotting the responses of the association curves against the DNA concentration and fitted using the following binding curve:

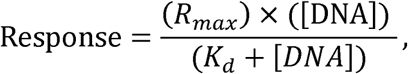

where R*_max_*refers to the maximum response due to the DNA binding to the sensor-immoblized protein.

### Structural modeling

All AlphaFold models were generated from the predicted AlphaFold Protein Structure Database^103^ or modeled *de novo* using the AlphaFold3 structure prediction server.^104^ To generate the intact WT model bound to the PAO1 *feo* upstream region, two copies of the respective protein sequence was used as input into the AlphaFold3 structure prediction server along with the following DNA duplex sequence: 5’-TCCACTCCGATACTGCGTCCTTAAGCGAGCCTTAAGTGAGAGTCGTTTAGATTCTC-3’ and 5’-GAGAATCTAAACGACTCTCACTTAAGGCTCGCTTAAGGACGCAGTATCGGAGTGGA-3’. The pLDDT score and Predicted aligned error (PAE) for this structure is provided in Fig. S6. AlphaFold3 structural predictions were carried out without any modifications of the input parameters.

### RNA sequencing

Total RNA was collected from mid-log phase WT PAO1 and Δ*bqsR* cells grown in BMM using the RNeasy Mini Kit (Qiagen). Purified RNA was treated with Turbo DNase (Invitrogen) and contaminating DNA was assessed using PCR with primers specific for the 16S rRNA gene (Table S6). RNA concentrations were determined using a NanoDrop spectrophotometer (Thermo Fisher Scientific) and the RNA quality was analyzed using an RNA 6000 Nano Chip on a Bioanalyzer 2100 (Agilent) and by 1.2 % agarose gel electrophoresis. RNA samples with A_260_/A_280_ ratio of 1.8-2.0 and an RIN value ≥ 7.0 were selected for sequencing analysis. Ribosomal RNA was removed using the Ribo-Zero rRNA Depletion Kit (Illumina). For sequencing, 250-300 bp insert size, strand-specific libraries were prepared and sequenced by Novogene Co., Ltd using paired-end sequencing (2 x 150 bp) on a Nova Seq 6000 platform.

### Analysis of RNA-seq reads

All raw RNA sequences were further analyzed using Kbase (https://www.kbase.us/). Three biological replicates were used for each stain. Reads were checked for quality using FastQC-v0.11.5 and trimmed using Trimmomatic-v0.36. The trimmed reads were aligned to the PAO1 genome using HISAT2-v2.1.0 and assembled using Cufflinks-v2.2.1. A differential expression matrix was generated using DESeq2-v1.20.0, and the resulting raw gene count matrix was uploaded to iDEP2.01 (http://bioinformatics.sdstate.edu/idep/) to determine log_2_ of fold change (FC) in gene expression in Δ*bqsR* compared to WT along with their false discovery rate (adjusted *p*-values). A volcano plot depicting differentially expressed genes (log_2_FC > |1|) with significant –log10 *p*-values (> 1.3 for *p* < 0.05) was generated using Rstudio. The differentially expressed genes were sorted into functional categories using annotations from Gene Ontology (https://www.geneontology.org/), Kyoto Encyclopedia of Genes and Genomes (KEGG) (https://www.kegg.jp/kegg/pathway.html), and the Pseudomonas Community Annotation Project (PseudoCAP) (https://pseudomonas.com/). Heatmaps of selected pathways were generated using MORPHEUS (https://software.broadinstitute.org/morpheus/).

### Reverse transcription-quantitative PCR (RT-qPCR)

Cultures were grown and RNA was collected following the same procedures outlined in the RNA sequencing methods. Extracted RNA was used as a template to synthesize cDNA using the Transcriptor First Strand cDNA Kit (Roche). Generation of cDNA was validated by PCR using primers specific for the 16S rRNA gene (Table S6). Real-time PCR amplifications using the LightCycler 480 SYBR Green I Master mix (Roche) were carried out in a LightCycler 480 II (Roche) using cDNA as the templates and primers specific for the house-keeping genes, *nadB* (PA0761) and *rpoD* (PA0576), and experimental genes (Table S6). Primer specificity was evaluated by using melting curve analysis with a single peak indicating the amplification of a single product. The transcript abundance was quantified using the standard curve method^105^ and normalized by the transcripts of house-keeping genes *rpoD* and *nadB*. Experiments were performed using three independent biological replicates per strain. To calculate the log_2_ fold changes (LFC) of gene expression between Δ*bqsR* and the WT, the transcript number derived from the Cp values from each individual Δ*bqsR* replicate was normalized by the average of the Cp-derived transcript numbers of three WT biological replicates. To assess the statistical significance and reproducibility, 95% confidence intervals were calculated for each gene. Ratios were determined by comparing the mean transcript number (n = 3 biological replicates) in Δ*bqsR* strain to that in the WT strain. Results were considered statistically significant if the 95 % confidence interval (CI) for the fold-change did not include 1. Using this approach, the transcriptional regulation of all examined genes was deemed significant.

### Generation of the *feo* promoter (P*^feo^*) fusion reporters

For plasmid-based reporter constructs, first the *feo* promoter was amplified and ligated into the digested pMS402 plasmid directly upstream of the *luxCDABE* operon. The ligated product was transformed into heat-shock competent *E. coli* DH5α cells. Successful transformants were selected on LB agar plates containing 50 µg/mL Kan. The resulting constructs were verified to have the insert by Nanopore DNA sequencing performed by Plasmidsaurus and are hereafter referred to as pAHT001. The pAHT001 construct was electroporated into the WT and Δ*bqsR* strains. The successful transformants were selected on LB agar plates containing 300 µg/mL Tmp.

### Growth studies and promoter assay

First, the level of ascorbic acid (AA) was optimized to limit Fe^2+^ oxidation without causing an inhibitory impact on *P. aeruginosa* growth. For this, pre-cultures were started from streaked colonies on LB plates in BMM8.5 with no added FeSO_4_ and incubated until mid-log (12 h) at 37 °C shaking at 200 rpm. The pre-cultures were normalized to OD_600_ of 0.3, inoculated 1:100 into fresh BMM8.5 containing 3.6 µM FeSO_4_ and aliquoted (200 µL) into the wells of a clear 96-well plate. AA stocks were made in increasing concentrations so that the addition of 5 µL of the stocks achieved final concentrations of 1.0, 1.5, 2.0, 2.5, and 3.0 mM. Water was added as a no-AA control. The growth of the cultures was monitored by measuring OD_600_ hourly in a BioTek SynergyMx until stationary phase (24 h) at 37 °C under medium shaking. Since addition of AA at 1.5 and 2.0 mM had the least inhibition of growth, these concentrations were considered for preventing oxidation of Fe^2+^. For this, the levels of Fe^2+^ were monitored using a well-established ferrozine-based assay^107^ with modifications. Briefly, 75 µL aliquots from each sample were transferred to a well of a clear 96-well plate. To the non-reduced samples, 14 µL of BMM8.5 with no iron was added; and 14 µL of AA (final 100 mM) was added to the reduced samples. Following 30 min incubation at room temperature, 18 µL of the ferrozine reagent was added (final 0.5 mM), incubated for 10 min, and the absorbance at 562 nm was measured. To set up the experiment, 10 mM FeSO_4_ stocks with increasing levels of AA were prepared so that the addition of 50 µL of the stocks to 4.95 mL of BMM8.5 with no iron resulted in the final concentration of 100 µM FeSO_4_ with 1.0, 1.5, and 2.0 mM AA. An aliquot of each medium (75 µL) was taken for an initial Fe^2+^ measurement and then again following static incubation for 6 h and 24 h at 37 °C to mimic the growth conditions. At 2.0 mM AA, the highest level of Fe^2+^ (20 µM) remained in the medium after 6 h and 24 h of static incubation. These conditions were used for promoter-reporter assays. We also evaluated the impact of pyruvate with reported antioxidant properties^108,109^ on mitigating the inhibitory effects of Fenton reactions likely stimulated by AA and Fe^2+^. For this, we monitored the growth of WT PAO1 cultured with 2.0 mM AA and 100 µM FeSO_4_ in the absence and presence of 10 mM pyruvate as described above. After 9 h of growth (mid-exponential phase), the cultures were treated with either 2 mM AA only or 4 µM and 100 µM FeSO_4_ with 2.0 mM AA. The cultures were incubated for an additional 24 h, measuring OD_600_ hourly. The addition of 10 mM pyruvate abrogated growth inhibition observed at 100 µM FeSO_4_ and was therefore used for promoter-reporter studies designed as follows: *P. aeruginosa* strains were grown in no iron-BMM8.5 with 10 mM pyruvate at 37 until mid-log phase (12 h), normalized to an OD_600_ of 0.3, and inoculated 1:100 into fresh no iron-BMM8.5 with 10 mM pyruvate. Cultures were grown statically (to minimize oxidation of Fe^2+^) in clear 96-well plates at 37 in a BioTek SynergyMx. The absorbance at 600 nm was measured hourly. After reaching mid-log phase (12 h), the cultures were treated with freshly prepared FeSO_4_ solutions in AA to achieve a final concentration of 4 µM or 100 µM FeSO_4_ and 2.0 mM AA. Treated cultures were monitored for luminescence and OD_600_ measured at hourly intervals. Relative luminescence units (RLU) were calculated after subtracting non-inoculated controls and normalizing by OD_600_. To report changes in P*^feo^*-dependent luminescence, log_2_(RLU_Δ*bqsR*_ /RLU*_WT_*) was calculated. Every experiment contained at least three independent biological replicates and was repeated at least twice for consistency. Statistical significance was evaluated by using 95 % confidence intervals.

### Metal-binding assays

(His)_6_-cleaved *Pa*BqsR constructs (intact WT, D51E, or DBD) were tested for their ability to bind Fe^2+^. Briefly, each construct was degassed and brought into an anoxic chamber containing an N_2_/H_2_ atmosphere and operating at < 5 ppm O_2_. After overnight equilibration with the N_2_/H_2_ atmosphere, each protein solution was incubated with 1-5 mol eq. of Fe^2+^ to assess stability of the protein in the presence of metal. The intact protein only tolerated a maximal loading of 1.2 mol eq. of Fe^2+^, so all metal-loading experiments were performed with this slight excess of metal cation in solution. After the addition of metal and incubation for 10 min at *ca.* 6 °C, the protein-metal solution was buffer exchanged into SEC buffer (50 mM Tris, pH 7.5, 100 mM NaCl, 5 % (v/v) glycerol, 1 mM TCEP) through extensive rounds of concentration and dilution using a 0.5 mL 10 kDa MWCO spin concentrator. A modified version of the ferrozine assay^110,111^ was performed as previously described.^112^ All metal loading was then further confirmed by measuring metal content using a PerkinElmer NexION 300D mass spectrometer connected to inductively couple plasmon mass spectrometry (ICP-MS).

### Circular dichroism spectroscopy

Circular dichroism (CD) spectra were collected using a JASCO J-710 spectropolarimeter. Samples at 0.2 mg/mL protein concentration in SEC buffer were contained within a 1 mm quartz cuvette. Plotted CD spectra are the average of five scans that were measured from 300 nm to 195 nm with a 0.1 nm data interval and a 5 s data averaging time.

### X– ray absorption spectroscopy (XAS)

**X-** Intact WT *Pa*BqsR samples were metal-loaded anoxically and concentrated to *ca.* 2 mM Fe^2+^ (final concentration), mixed anoxically with ethylene glycol (20% (v/v) final), and were aliquoted anoxically into Lucite cells wrapped with

Mylar tape, flash-frozen in liquid N_2_ and stored at –80 °C until data collection. X-ray absorption spectroscopy (XAS) was performed on beamlines 7-3 and 9-3 at the Stanford Synchrotron Radiation Lightsource (Menlo Park, CA), and samples in replicates were collected when possible. Extended X-ray absorption fine structure (EXAFS) of Fe (7210 eV) was measured using a Si 220 monochromator with crystal orientation φ = 90°. Samples were measured as frozen aqueous glasses at 15 K, and the X-ray absorbance was detected as Kα fluorescence using either a 100-element (beamline 9-3) or 30-element (beamline 7-3) Ge array detector. A Z-1 metal oxide filter (Mn) and slit assembly were placed in front of the detector to attenuate the elastic scatter peak. A sample-appropriate number of scans of a buffer blank were measured at the absorption edge and subtracted from the raw data to produce a flat pre-edge and eliminate residual Mn Kβ fluorescence of the metal oxide filter. Energy calibration was achieved by placing a Fe metal foil between the second and third ionization chamber. Data reduction and background subtraction were performed using EXAFSPAK (Microsoft Windows version).^113^ The data from each detector channel were inspected for drop outs and glitches before being included into the final average. EXAFS simulation was carried out using the program EXCURVE (version 9.2) as previously described.^113–115^ The error associated with the XAS data are as follows: distances, ± 0.02 Å; coordination number, ± 25 %; identity of scattering atom, ΔZ ± 1 (Z=6-17), ΔZ ± 3 (Z=20-35).^116^ The quality of the fits was determined using the least-squares fitting parameter, *F*, which referred to as the fit index (FI) and is defined as:

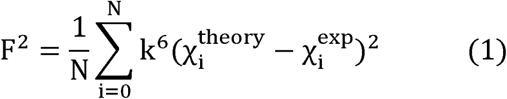

## Data Availability

Sourced data are provided with this paper. The data generated in this study are provided in the Supplementary Information/Source Data file. The following Uniprot IDs were used in this study: *Pseudomonas aeruginosa* strain PAO1 BqsR, <u>Q9I0I1</u>; *Escherichia coli* strain K12 KdpE, <u>P21866</u>; *Escherichia coli* strain K12 OmpR, <u>P0AA16</u>; *Escherichia coli* strain K12 PhoB, <u>P0AFJ5</u>; *Klebsiella pneumoniae* strain ATCC 700821 / MGH 78578 RstA, <u>A6T8N1</u>. All unique biological materials (*e.g.*, expression plasmids, bacterial strains) are readily available from the authors upon request.

## Abbreviations

AA, ascorbic acid; ANR, anaerobic transcriptional regulator; BLI, biolayer interferometry; CD, circular dichroism; CF, cystic fibrosis; CI, confidence interval; Cp, crossing point; CV, column volumes; DBD, DNA-binding domain of *Pa*BqsR; DEGs, differentially expressed genes; EMSA, electrophoretic mobility shift assays; EXAFS, extended X-ray absorption fine structure; FI, fit index; FUR, ferric uptake regulator; IMAC, immobilized metal affinity chromatography; ICP-MS, inductively couple plasmon mass spectrometry; IPTG, isopropyl β-D-1-thiogalactopyranoside; LB, Luria broth; LFC, log_2_fold change; HK, membrane sensor histidine kinase; MWCO, molecular weight cutoff; RD, N-terminal receiver domain; NMR, nuclear magnetic resonance; NOESY, nuclear Overhauser effect spectroscopy; OD, optical density; *Pa*BqsR, the response regulator *Pseudomonas aeruginosa* BqsR; *Pa*BqsS, the histidine kinase *Pseudomonas aeruginosa* BqsS; PMSF, phenylmethylsulfonyl fluoride; PAE, predicted aligned error; RLU, relative luminescence units; RNA-seq, RNA sequencing; RR, response regulator; SEC, size-exclusion chromatography; TEV, tobacco etch virus; TCS, two-component signal transduction system; UV, ultraviolet; WT, wildtype; XANES, X-ray absorption near-edge structure; XAS, X-ray absorption spectroscopy.

## Acknowledgements

This work was supported by NIH-NIGMS grant R35 GM133497 (A.T.S), NIH-NIGMS grant T32 GM158458 (A. T. S. and A. P.), HHMI Gilliam Fellowship GT15765 (A. P.), NIH-NIGMS grant R15 GM124670 (M.A.P.), grant CFF-005124P222 (M.A.P.), and the Murdock Trust Swanson Promise for Scientific Research Award (K.N.C.). Use of the Stanford Synchrotron Radiation Lightsource, SLAC National Accelerator Laboratory, is supported by the U.S. Department of Energy, Office of Science, Office of Basic Energy Sciences under Contract No. DE-AC02-76SF00515. The SSRL Structural Molecular Biology Program is supported by the DOE Office of Biological and Environmental Research, and by the National Institutes of Health, National Institute of General Medical Sciences (P30GM133894). The contents of this publication are solely the responsibility of the authors and do not necessarily represent the official views of NIGMS or NIH. Sequence searches utilized both database and analysis functions of the Universal Protein Resource (UniProt) Knowledgebase and Reference Clusters (http://www.uniprot.org) and the National Center for Biotechnology Information (http://www.ncbi.nlm.nih.gov/). We thank Prof. Michael F. Summers and Dr. Nele Hollmann for both the usage of their NMR facilities and their extensive help with NMR data analysis. We thank Dr. Chelsea Murphy at the Oklahoma State University Bioinformatic Center for her assistance with RNA-seq plots. We also thank Jacob Burch-Konda for the initial analysis of the RNA sequencing data and Alexis J. Northrup for the initial growth studies.

## Author contributions

A.P., M.H., H.S., J.B.B., K.N.C., M.A.P., and A.T.S. designed the research; A.P., M.H., H.S., D.G., A.O.T., A.K.; S.P.; R.K.; J.B.B., and K.N.C. performed the research; A.P, M.H., H.S., K.N.C., M.A.P., and A.T.S. analyzed the data; A.P., M.H., K.N.C., M.A.P., and A.T.S. wrote and edited the paper.

## Additional information

Supplementary information. Supplementary information accompanies this paper that includes supplementary figures S1-S16, supplementary tables S1-S6, as well as any additional experimental details, materials, and methods.

## Figure Legends

