## Supporting Information for "The response regulator BqsR/CarR controls ferrous iron (Fe^2+^) acquisition in *Pseudomonas aeruginosa*"

**Table S1**. *Pa*BqsR N-terminal regulatory domain (residues 1-123; PDB ID 8GC6) data collection and refinement statistics. Parentheses indicate the highest resolution shell.

| | |
|---|---|
| **Data Collection** | |
| Beamline | APS 21-ID-F |
| Wavelength (Å) | 0.97872 |
| Space group | $P4_12_12$ |
| Cell Dimensions | |
| *a*, *b*, *c* (Å) | 46.74, 46.74, 111.16 |
| α, β, γ (˚) | 90, 90, 90 |
| Resolution (Å) | 29.04-1.30 (1.32-1.30) |
| $R_{merge}$ | 0.047 (1.263) |
| $I/\sigma(I)$ | 17.7 (0.9) |
| Completeness (%) | 99.41 (94.63) |
| **Refinement** | |
| Resolution (Å) | 29.04-1.30 (1.32-1.30) |
| No. reflections | 31032 |
| $R_{work}$ | 0.158 |
| $R_{free}$ | 0.175 |
| No. atoms | |
| Protein | 950 |
| Water | 106 |
| Average B-factors (Å$^2$) | 21.00 |
| R.m.s. deviations | |
| Bond lengths (Å) | 0.013 |
| Bond angles (˚) | 1.48 |
| Ramachandran plot | |
| Favored | 100 % |
| Allowed | 0 % |
| Outlier | 0 % |

**Table S2**. Structural restraints and refinement statistics for the NMR structure of the *Pa*BqsR C-terminal DNA-binding domain (residues 123-223; PDB ID 9YFA and BMRB ID 31268).

| **NMR-derived restraints** | |
|---|---|
| Distance restraints | |
| Total | 1912 |
| Intraresidue & interresidue sequential, \|i-j\|=0,1 | 628 |
| Interresidue short-range \|i-j\|<=1 | 628 |
| Interresidue medium-range, 1<\|i-j\|<5 | 382 |
| Interresidue long-range, \|i-j\|>=5 | 274 |
| Hydrogen bonds | 48 |
| Dihedral $\phi$ and $\psi$ angle constraints from TALOS | 148 |
| CYANA structural statistics | |
| RMSD to mean structure (Å): | |
| Backbone | 0.93 Å ± 0.11 Å |
| Heavy atom | 1.43 Å ± 0.10 Å |
| Ramachandran analysis | |
| Most favored regions | 75.9 % |
| Additional allowed regions | 23.4 % |
| Generously allowed regions | 0.7 % |
| Disallowed regions | 0.0 % |

**Table S3**. Examples of select PAO1 gene upstream regions related to biofilm formation/dispersal and/or metal homeostasis that contain the BqsR box consensus sequence.

| Consensus sequence: 5'-TTAAGNNNNNNTTAAG-3' | |
|---|---|
| **Upstream sequence** | **Gene name** |
| 5'- N…TTAAGCGAGCCTTAAG…N-3' | *feoABC* ($Fe^{2+}$ transporter) |
| 5'- N…TTAAGCACCTCTTAAG…N-3' | *speD2* (biofilm formation regulator) |
| 5'-N…TTAAGGGACAATTAAG…N-3' | *PA0545* (ferric reductase) |
| 5'- N…TTAAGGAATCCTTAAGGTTTGCTTAAG…N-3' | *algD* (involved in biofilm formation) |

**Table S4**. List of primers and DNA sequences used to make variant proteins, for EMSAS, and for DNA analysis in this study. Underlines indicate the modified bases for the introductions of mutations.

| Name | Description | Oligonucleotide sequence |
|---|---|---|
| D51E_Fwd | Used for D51E site-directed mutagenesis mutation | 5'-CCCGGCAGACCCAGTTCCAGAATGATCAG-3' |
| D51E_Rev | Used for D51E site-directed mutagenesis mutation | 5'-CTGATCATTCTGGAACTGGGTCTGCCGGG-3' |
| R211A_Fwd | Used for R211A site-directed mutagenesis mutation | 5'-CGAAACCCGTGCTGGTCAAGGTTAC-3' |
| R211A_Rev | Used for R211A site-directed mutagenesis mutation | 5'-GTAACCTTGACCAGCACGGGTTTCG-3' |
| R211K_Fwd | Used for R211K site-directed mutagenesis mutation | 5'-CGAAACCCGTAAAGGTCAAGGTTAC-3' |
| R211K_Rev | Used for R211K site-directed mutagenesis mutation | 5'-GTAACCTTGACCTTTACGGGTTTCG-3' |
| *feo*_upstream_21bp | Double-stranded DNA used in EMSAs | 5'-CCTTAAGCGAGCCTTAAGTGA-3' and 5'-TCACTTAAGGCTCGCTTAAGG-3' |
| *carO*_upstream_26bp | Double-Stranded DNA used in EMSAs | 5'-GCCGATTAAGCCTGGTTTCAGCAAGG-3' and 5'-CCTTGCTGAAACCAGGCTTAATCGGC-3' |
| *feo*_upstream_56bp | Double-stranded DNA sequence used for larger DNA EMSAs and for AlphaFold modeling | 5'-TCCACTCCGATACTGCGTCCTTAAGCGAGCCTTAAGTGAGAGTCGTTTAGATTCTC-3' and 5'-GAGAATCTAAACGACTCTCACTTAAGGCTCGCTTAAGGACGCAGTATCGGAGTGGA-3' |
| DNA_test_16S_rRNA_Fwd | Used in PCR test to assess for contaminating DNA | 5'-GTGCCAGCAGCCGCGGTAA-3' |
| DNA_test_16S_rRNA_Rev | Used in PCR test to assess for contaminating DNA | 5'-AGGGTTGCGCTCGTTG-3'. |
| Scrambled_*feo*_upstream | Scrambled double-stranded DNA sequence used for EMSAs | 5'-GCCTCTGCCGAGCCTATCCTGA-3' and 5'-TCAGGATAGGCTCGGCAGAGGC-3' |

**Table S5**. Fits obtained for the Fe K-edge EXAFS of WT *Pa*BqsR titrated with $Fe^{2+}$. Fits were determined by curve fitting using the program EXCURVE (version 9.2).

| | | Fe-N (His) | | | Fe-O/N (non His) | | | $E_o$[e] (eV) |
|---|---|---|---|---|---|---|---|---|
| **Sample** | **Fit index**[a] | **No**[b] | **R**[c] **(Å)** | **DW**[d] **(Å$^2$)** | **No** | **R (Å)** | **DW (Å$^2$)** | |
| WT *Pa*BqsR + $Fe^{2+}$ | 0.085 | 4[f] | 2.09[f] | 0.002 | 2[f] | 2.17[f] | 0.007 | -1.96 |

[a]The least-squares fitting parameter (see *Methods*) [b]Coordination number [c]Bond length [d]Debye-Waller factor [e]Photoelectron energy threshold [f]The error associated with the XAS data are as follows: distances, ± 0.02 Å; coordination number, ± 25 %; identity of scattering atom, ΔZ ± 1 (Z=6-17), ΔZ ± 3 (Z=20-35)

**Table S6**. Bacterial strains, plasmids, and primers used for *in vivo* experiments in this study.

| Strain/plasmid/primers | Description | Reference |
|---|---|---|
| ***P. aeruginosa* strains** | | |
| PAO1 | Laboratory wild type of *P. aeruginosa* PAO1 | 1 |
| Δ*bqsR* | *bqsR* deletion in PAO1 background | 2 |
| PAO1/pAHT001 | PAO1 harboring plasmid-based *feo* promoter reporter | This study |
| Δ*bqsR*/pAHT001 | Δ*bqsR* harboring plasmid-based *feo* promoter reporter | This study |
| ***E. coli* strains** | | |
| DH5α | General purpose cloning strain | NEB |
| **Plasmids** | | |
| pMS402 | *P. aeruginosa* promoter-less vector harboring *luxCDABE*, $Tmp^R$ | 3 |
| pAHT001 | pMS402 harboring 300 bp of *feo* promoter region ligated using XhoI and BamHI, $Tmp^R$ | This study |
| **Primers** | | |
| XhoI_feo_pro_F | 5'-TTTCTCGAGCGCCTTCCGCACCCAG-3' | This study |
| BamHI_feo_pro_R | 5'-TTTGGATCCGGCGGATTCCAGGCAGAGG-3' | This study |
| 16S_533_F | 5'-GTGCCAGCAGCCGCGGTAA-3' | 4 |
| 16S_1100_R | 5'-AGGGTTGCGCTCGTTG-3' | 4 |
| nadB_914_F | 5'-GGATCGACTGCGTCTACCTG-3' | This study |
| nadB_1016_R | 5'-ATGTCGATGCCGAAGTCCAG-3' | This study |
| rpoD_F | 5'-GGGCGAAGAAGGAAATGGTC-3' | This study |
| rpoD_R | 5'-CAGGTGGCGTAGGTGGAGAA-3' | This study |
| carO_159_F | 5'-CATCGTCAAGCGCATCAAGGG-3' | This study |
| carO_264_R | 5'- GCGTTCCACAAGTTCAGCCC -3' | This study |
| speD2_163_F | 5'- ATGCTCGAGATCAGGTGCCG -3' | This study |
| SpeD2_359_R | 5'-GTACGTCGCCGCAGGTGAA-3' | This study |
| lptF_PA3692_70_F | 5'-CCGAACGCCAACCTGGAACA-3' | This study |
| lptF_PA3692_262_R | 5'-GTTCGATGCGCTGGTTGGTC-3' | This study |
| feoB_2005_F | 5'- GTCCTGCTCTACGTGCCCTG-3' | This study |
| feoB_2178_R | 5' GATGGTCAGGACGCTACGCT-3' | This study |
| opdQ_649_F | 5'- GGCAGCTACAAGTGGACCGA-3' | This study |
| opdQ_841_R | 5'- CGTTGAGCGCCTTGTTGTCG-3' | This study |
| bqsS_225_F | 5'-GTATTCCGGGCGCTATTTCG-3' | This study |
| bqsS_330_R | 5'-CTGCACCAGCCCTTTCTTG-3' | This study |

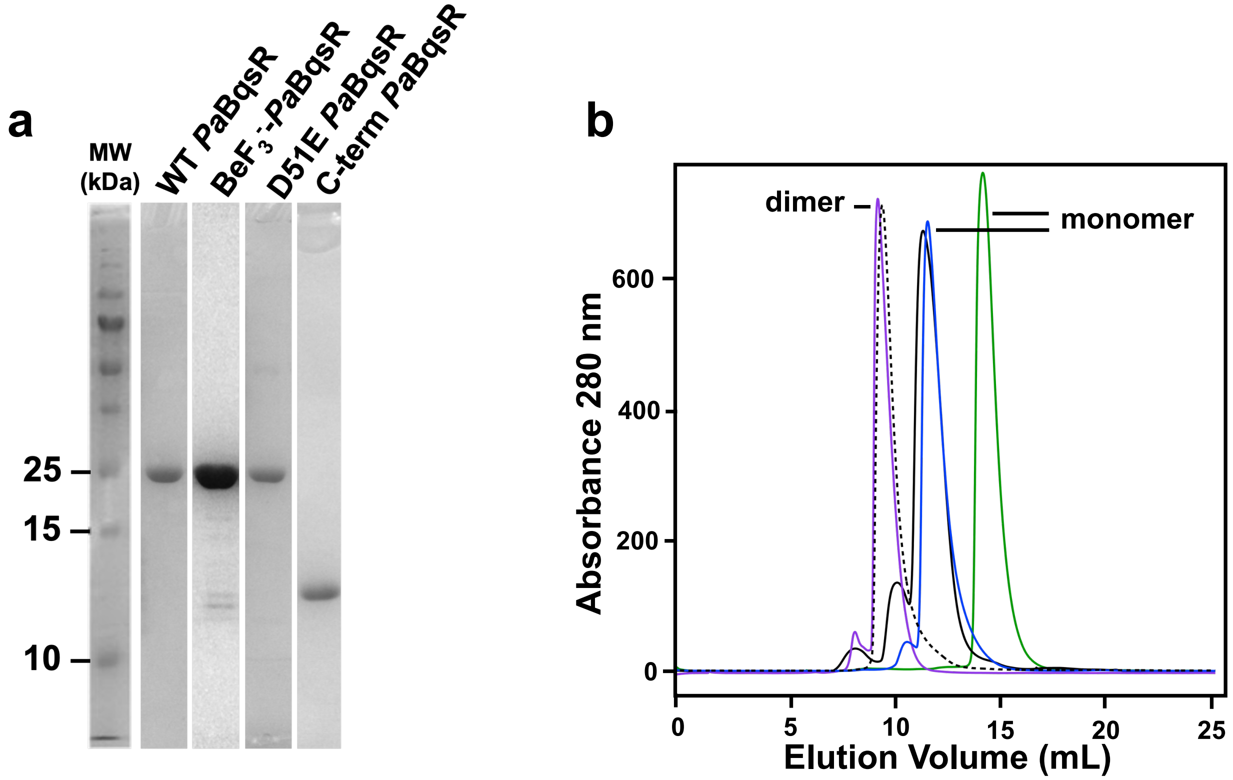


**Figure S1**. Purified *P. aeruginosa* BqsR. **a.** 15 % SDS-PAGE analysis of WT *Pa*BqsR, $BeF_3^-$-*Pa*BqsR, D51E *Pa*BqsR, and the C-term *Pa*BqsR DNA-binding domain (DBD). **b**. Size exclusion chromatography (SEC) profiles of low concentration (<< 1 mM) WT *Pa*BqsR (black), high concentration (≥ 1 mM) WT *Pa*BqsR (purple), D51E *Pa*BqsR (blue), the C-term *Pa*BqsR DNA-binding domain (DBD) (green), and $BeF_3^-$-activated *Pa*BqsR (dashed). Highly concentrated WT *Pa*BqsR (≥ 1 mM) and $BeF_3^-$-activated *Pa*BqsR both migrate as dimers, while low concentration (<< 1 mM) WT *Pa*BqsR, D51E *Pa*BqsR, and the C-term *Pa*BqsR DBD all migrate as monomers.

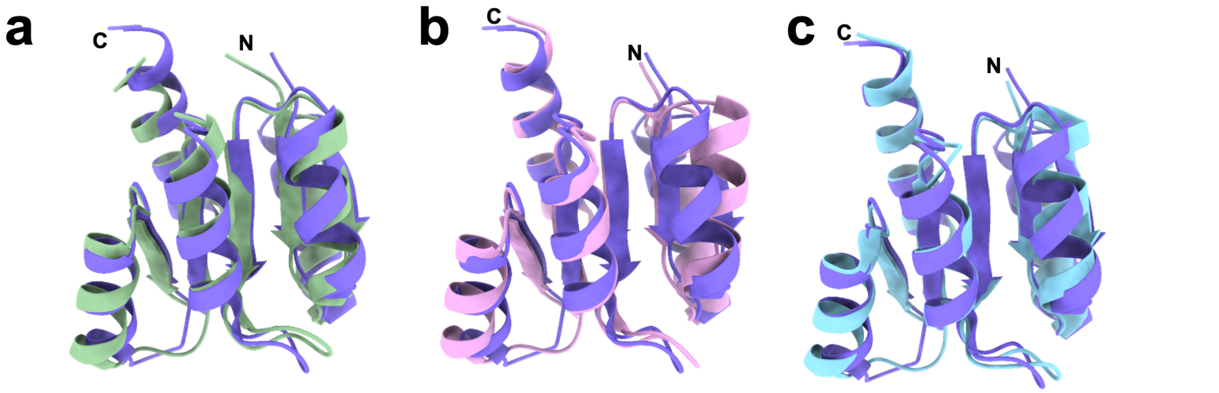


**Figure S2**. Superpositioning of the *Pa*BqsR N-terminal receiver domain (purple; PDB ID 8GC6) reveals strong structural homology to other members OmpR/PhoB RR family. **a**. Superpositioning of *Pa*BqsR onto ArlR (green; PDB ID 6IS1) reveals a Cα RMSD value of 0.992 Å across 106 pruned atom pairs. **b**. Superpositioning of *Pa*BqsR onto PhoP (pink; PDB ID 2PL1) reveals a Cα RMSD value of 0.961 Å across 105 pruned atom pairs. **c**. Superpositioning of *Pa*BqsR onto KdpE (blue; PDB ID 1ZH2) reveals a Cα RMSD value of 0.899 Å across 105 pruned atom pairs. In all cases, 'N' and 'C' represent the N- and C-termini, respectively.

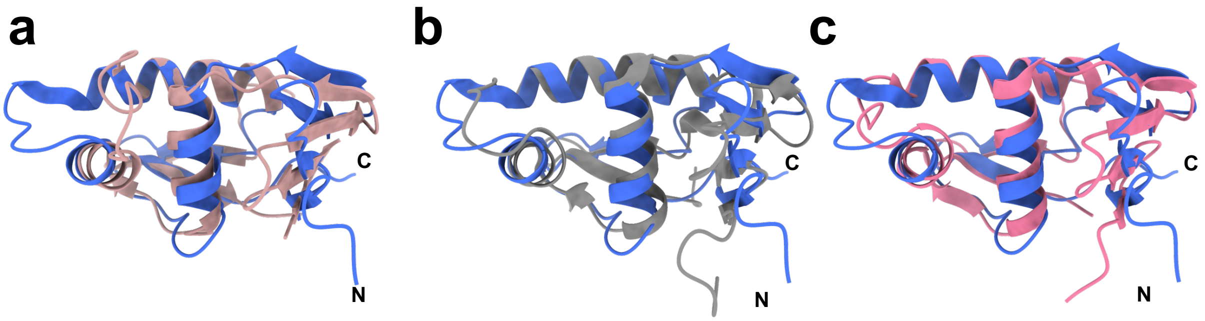


**Figure S3**. The NMR structure of the *Pa*BqsR DNA-binding domain (DBD) (blue) reveals a winged helix-turn-helix (HTH) motif, and superpositioning shows this motif to be common amongst the OmpR/PhoB RR family. **a**. Superpositioning of the *Pa*BqsR DBD with the OmpR DBD (brown; PDB ID 1OPC) reveals a Cα RMSD value of 1.13 Å across 103 pruned atom pairs. **b**. Superpositioning of the *Pa*BqsR DBD with the PhoP DBD (gray; PDB 1GXQ) reveals a Cα RMSD value of 1.12 Å across 103 pruned atom pairs. **c**. Superpositioning of the *Pa*BqsR DBD with the KdpE DBD (pink; PDB ID 3ZQ7) reveals a Cα RMSD value of 1.27 Å across 103 pruned atom pairs. In all cases, 'N' and 'C' represent the N- and C-termini, respectively.

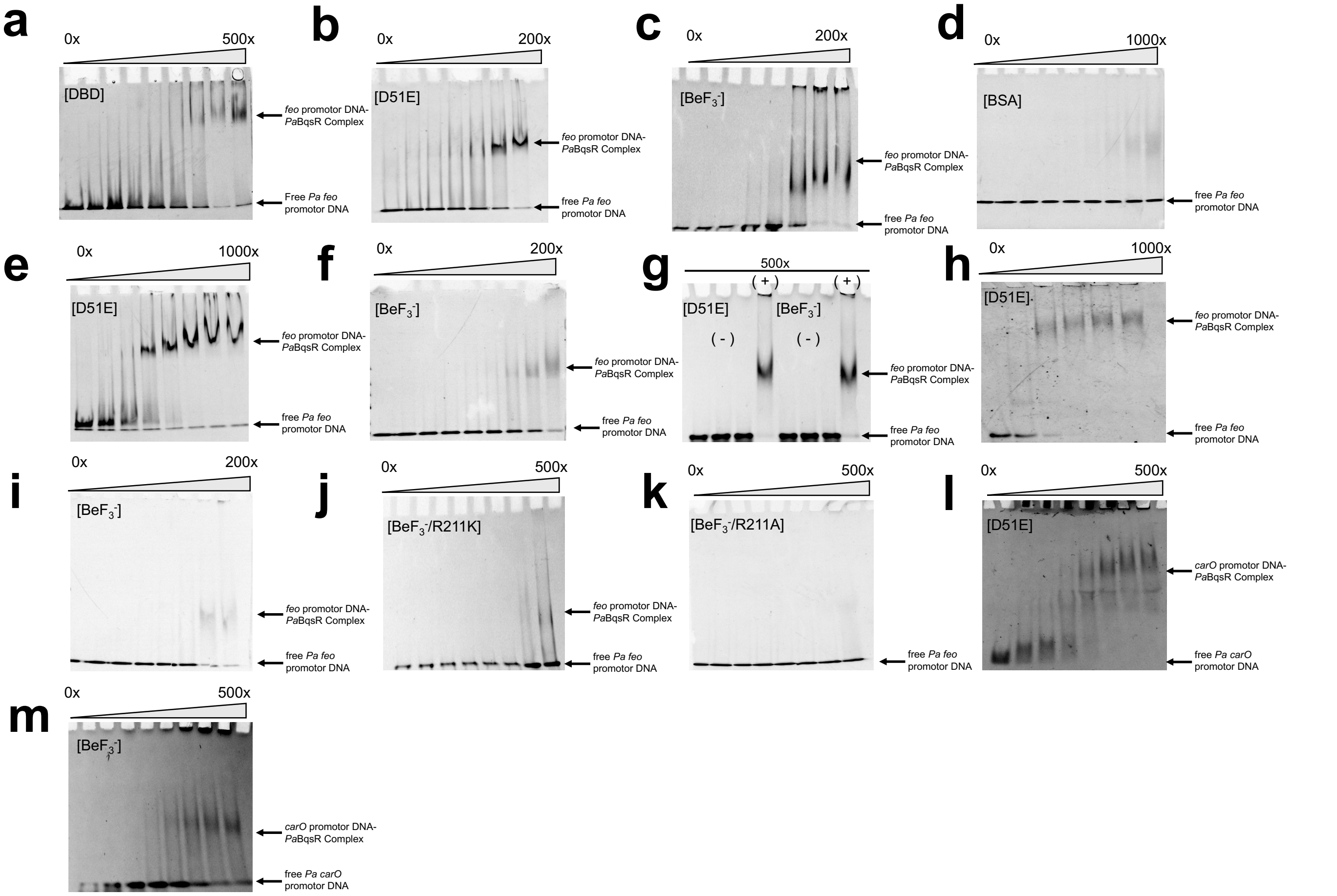


**Figure S4**. EMSAs of *Pa*BqsR in complex with the PAO1 *feo* upstream region reveal that the N-terminal phosphor-acceptor and dimerization domain increase affinity towards DNA. **a**. EMSAs reveal that the C-terminal *Pa*BqsR DBD only binds weakly to the PAO1 *feo* upstream region, requiring a *ca.* 500:1 ratio to reach binding saturation. **b**. EMSAs reveal that the pseudo-activated D51E *Pa*BqsR binds more strongly to the PAO1 *feo* upstream region, requiring only a *ca.* 200:1 ratio to reach binding saturation. **c**. EMSAs reveal that the phosphomimetic $BeF_3^-$-*Pa*BqsR binds to the PAO1 *feo* upstream region similarly as the D51E *Pa*BqsR variant and requires only a *ca.* 200:1 ratio to reach binding saturation. **d**. Control EMSA with the *feo* upstream region using bovine serum albumin (BSA) in place of *Pa*BqsR indicates no binding. The slight shading at 1000:1 ratio occurs due to the presence of BSA, while the band for the *feo* upstream region does not change in intensity. **e,f**. EMSAs using a larger, 56 bp oligonucleotide of the *feo* upstream region show that the approximate binding affinity is not dependent on the size of DNA. Both the D51E *Pa*BqsR variant (**e**) and the $BeF_3^-$-activated *Pa*BqsR (**f**) still require only a *ca.* 200:1 ratio to reach binding saturation. **g**. Control EMSAs with a scrambled *feo* oligonucleotide indicate *Pa*BqsR is selective for its palindromic sequence. (-) represents the scrambled oligonucleotide, while (+) represents the native *feo* oligonucleotide. All samples were run at 500x BqsR:DNA to ensure saturation. **h,i**. EMSAs performed using either D51E *Pa*BqsR (**h**) or $BeF_3^-$-activated *Pa*BqsR (**i**) at high salt concentration (500 mM NaCl) do not show a difference in binding behavior. **j,k**. EMSAs performed using either $BeF_3^-$-activated R211K *Pa*BqsR (**j**) or $BeF_3^-$-activated R211A *Pa*BqsR (**k**) at high salt concentration (500 mM NaCl) show decreased affinity, emphasizing that electrostatic interactions dominate *feo* binding for $BeF_3^-$-activated *Pa*BqsR in the absence of the key $Arg^{211}$ residue. **l,m**. EMSAs performed using either D51E (**l**) or $BeF_3^-$-activated *Pa*BqsR (**m**) reveal that although *carO* is negatively regulated in *bqsR* deletion studies, *Pa*BqsR binds to the BqsR box located on the *carO* promotor region, indicating that *Pa*BqsR can provide both positive and negative modes of regulation. All EMSAs were performed in triplicate with 0.5 μM DNA.

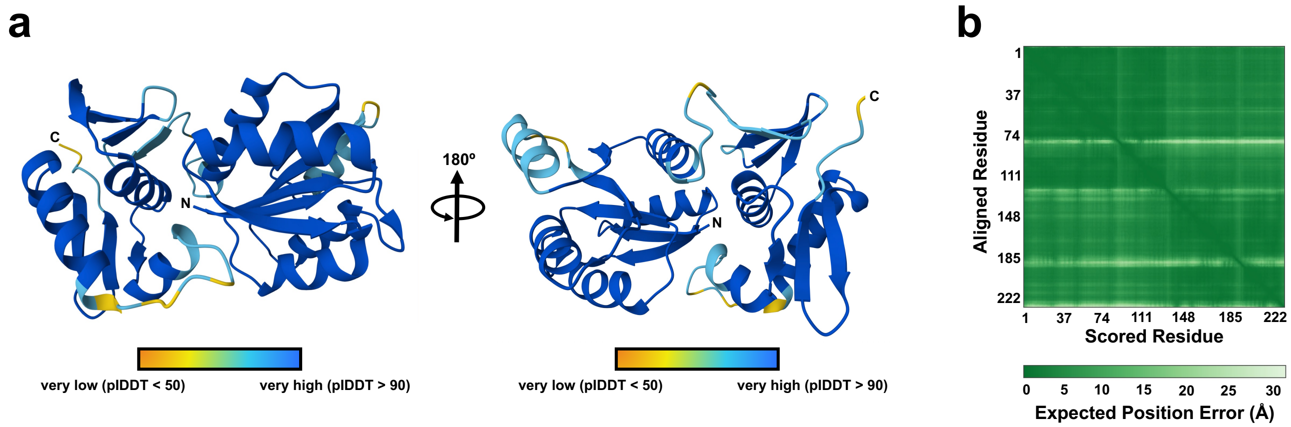


**Figure S5**. AlphaFold model of intact *Pa*BqsR color-coded based on the per-residue confidence score (pLDDT) from very low confidence (orange) to very high confidence (blue). **a**. pLDDT-colored AlphaFold model of *Pa*BqsR in two separate views rotated by 180°. **b**. Predicted aligned error (PAE) for the intact *Pa*BqsR AlphaFold model. In all cases, 'N' and 'C' represent the N- and C-termini, respectively.

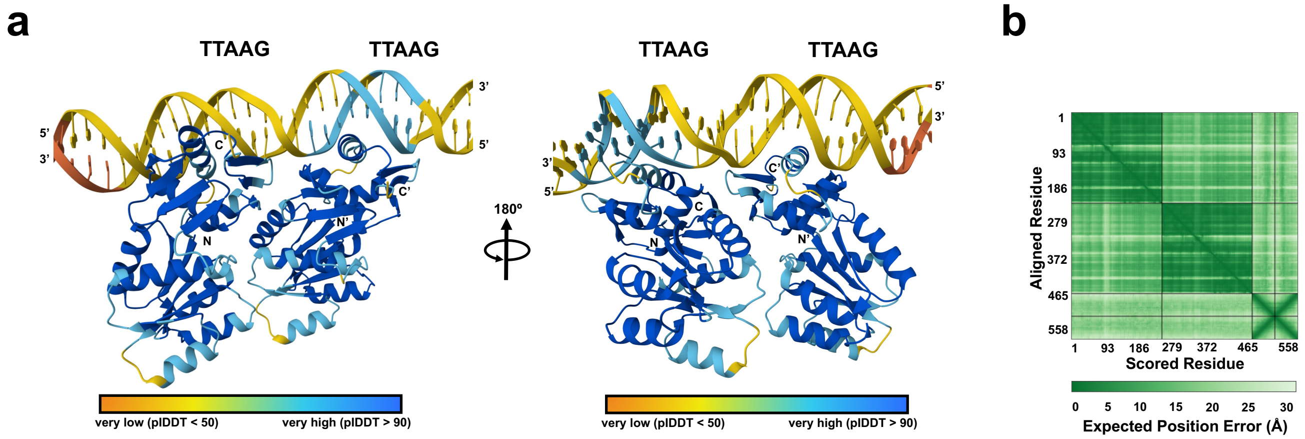


**Figure S6**. AlphaFold model of intact, dimeric *Pa*BqsR bound to the *feo* upstream region color-coded based on the per-residue confidence score (pLDDT) from very low confidence (orange) to very high confidence (blue). **a**. pLDDT-colored AlphaFold model of *Pa*BqsR bound to the *feo* upstream region in two separate views rotated by 180°. **b.** Predicted aligned error (PAE) for the intact, dimeric *Pa*BqsR bound to the *feo* upstream AlphaFold model. In all cases, 'N' and 'C' represent the N- and C-termini, respectively.

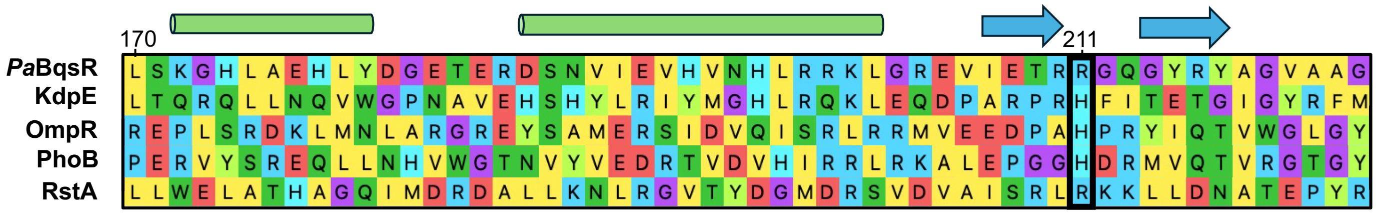


**Figure S7**. Partial multiple sequence alignment (MSA) of the DNA-binding domain (DBD) of various response regulators, such as *Pa*BqsR, *Ec*KdpE, *Ec*OmpR, *Ec*PhoB, and *Kp*RstA. Conserved structural elements such as the helix-turn-helix (HTH) motif (indicated by green cartoon cylinders, top) and a short β-hairpin (indicated by blue arrows, top) are present across these DBDs. However, despite structural conservation, sequence is not conserved in this structural element.

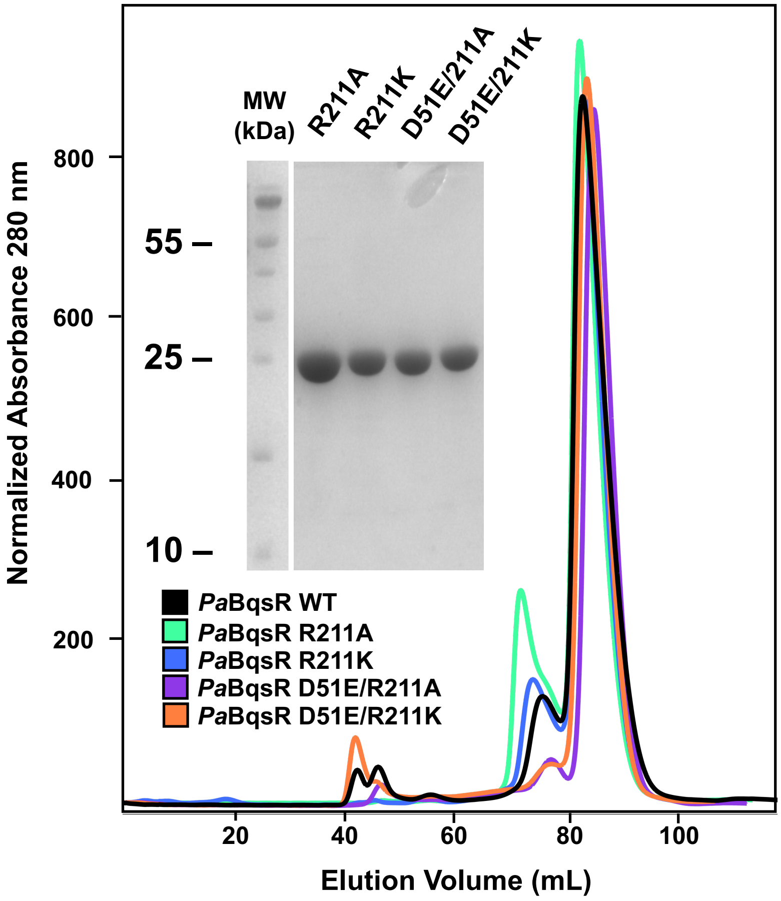


**Figure S8**. WT *Pa*BqsR and *Pa*BqsR variants R211A, R211K, D51E/R211A, and D51E/R211K have similar purity (*inset*: 15% SDS-PAGE), and all migrate similarly based on size-exclusion chromatography.

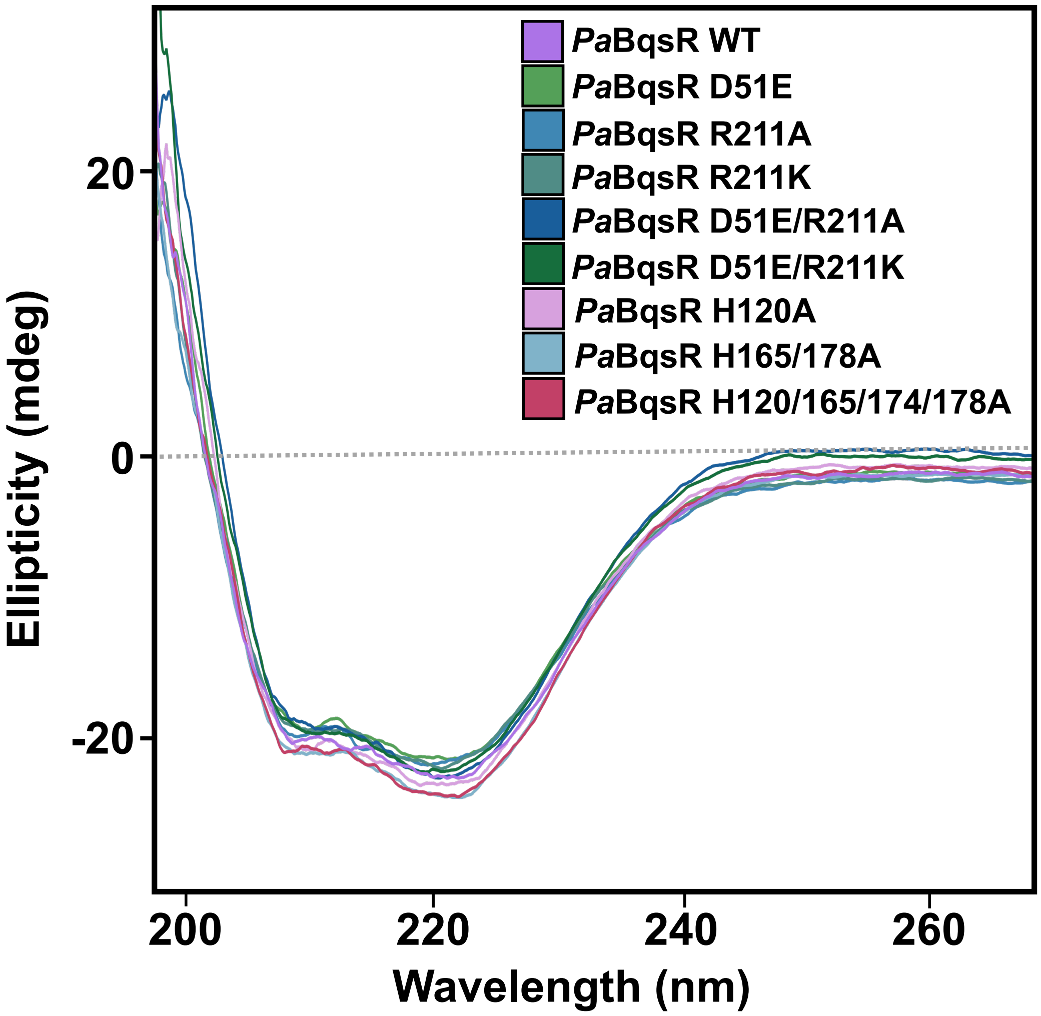


**Figure S9**. Circular dichroism spectra of all purified *P. aeruginosa* BqsR and variants used in this work. All variants exhibit nearly identical circular dichroism spectra compared to that of WT *Pa*BqsR, indicating that all of the variant proteins have gross secondary structures similar to the WT protein.

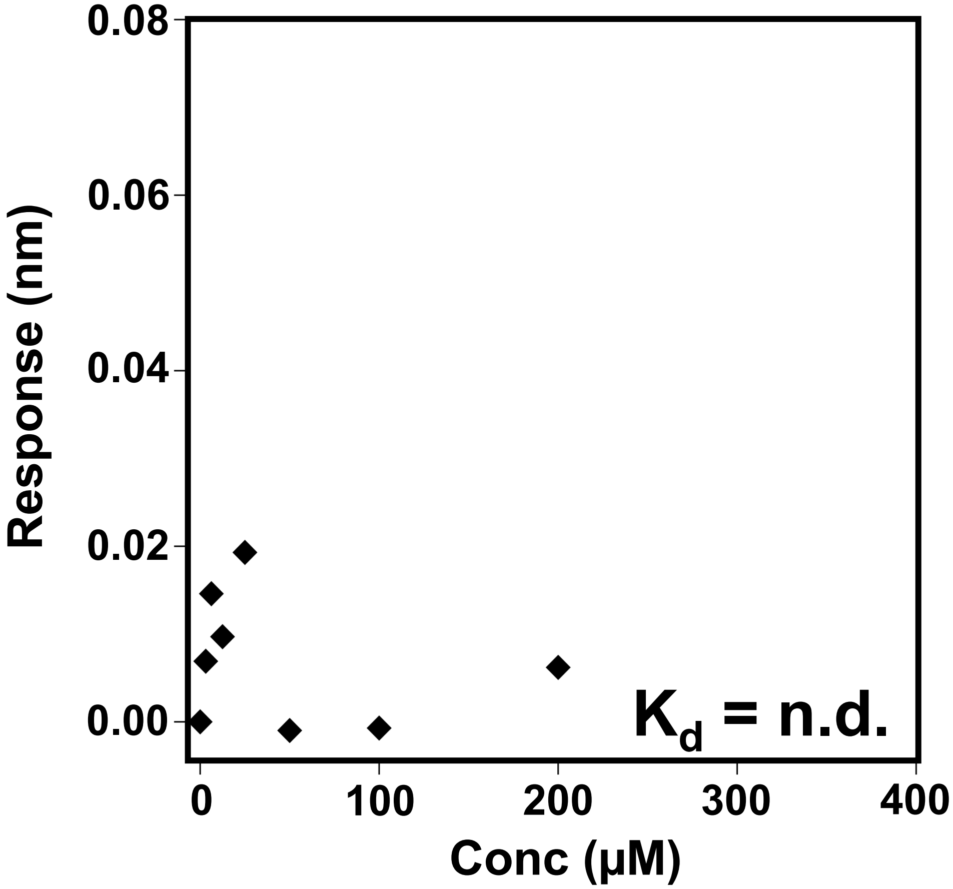


**Figure S10**. BLI data reveal that *Pa*BqsR has no measurable binding to the scrambled BqsR box (5'-GCCTCTGCCGAGCCTATCCTGA-3' and 5'-TCAGGATAGGCTCGGCAGAGGC-3'), emphasizing the specificity of *Pa*BqsR for its target DNA sequence.

| Gene name | BqsR Regulation: Fe2+ conc 4 µM | BqsR Regulation: Fe2+ conc 100 µM | Gene upstream sequence (5'→ 3') |
|---|---|---|---|
| *dhcR* (PA1998) | + | NA | **TTGTGA**TTGTGCGTGCGGCAGT**CTGACT**GCCGGGTCG<br>CACAGAGCTTCTCCGCTGCGGACTGATTTGTTAAGTTT<br>GTTTTTCC |
| ***carO* (PA0320)** | + | + | **TGGCCA**TCGACGACGAGGGCAC**CATCTA**CATGGTCAG<br>CGAGCCGAACCTGTTCTACGTGTTCCGCAAGAAGAGC<br>GGCGAGCGCCTGGCCACCAGCGACGCCGATTAAGCC<br>TGGTTTCAG |
| *carP* (PA0327) | + | + | **CGGACA**GGAAACTCGGACCTGT**CCGAGT**CTCCCGGGC<br>GGCTGGCTGACTTTAAGCTTGGCTTCAG |
| ***lptF* (PA3692)** | + | NA | **ATGCGC**GATACGCAGCACTAAGC**CGGATT**CCGCCCTTC<br>TTTCGTCGATTCGTGATTAAG |
| *rplQ* (PA4237) | - | NA | TTAAGGATGAA**TGACA**GCCCTCCTACAATCCA**ATTTAT** |
| *gltP* (PA5479) | - | NA | **TAGACG**GGGTAGGGGAGCGCATG**AATCAT**TGCGCCCT<br>TTCGCCTGTCGGGAGCGACTCTCCCGTTGGAGCGACT<br>CCCGACTTCCACAAGAATTAAG |
| ***opdQ* (PA3038)** | - | NA | **ATGTCT**AACAGTGGCCGTCTTC**CTGTAG**TGCTTAAG |
| ***feoB* (PA4358)** | - | + | **TGGATC**AGCGAATCCGCTGCTCC**TCCACT**CC**GATACT**<br>GCGTCCTTAAGCGAGCC**TTAAGT** |
| ***speD2* (PA4773)** | - | + | TTAAG**CACCTC**TTAAGTCAGCCGCGCC**TATCGT** |
| ***bqsS* (PA2656)** | - | + | **TTGCGG**CAAATTCAGCTTCAAT**TAAGAG** |

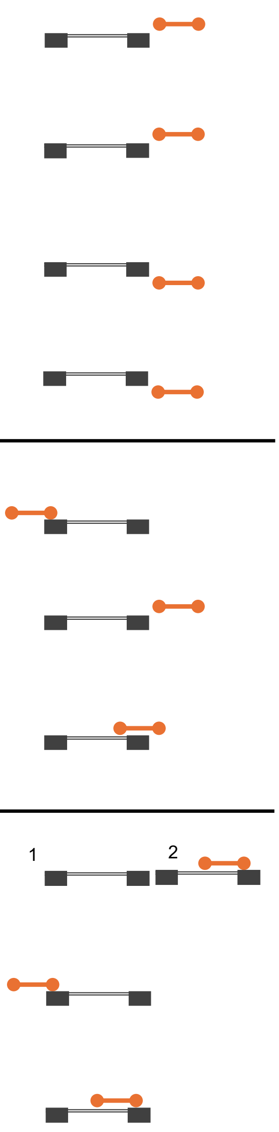


**Figure S11**. Placement of the BqsR box relative to the predicted -10/-35 regions of key gene promoters. A set of *Pa*BqsR-regulated genes with LFC ranging from -5.0 to 6.3 and possessing either the exact BqsR box (**5'-TTAAG**NNNNNN**TTAAG-3'**)[5] or its homolog (with deviating nucleotides shown in purple) detected using the MEME suite analysis were selected for the analysis. The -10/-35 regions (bolded in black) were predicted using BPROM (www.softberry.com/cgi-bin/programs/gfindb/bprom.pl) and DeNovoDNA (www.denovodna.com). Two -10/-35 regions were predicted for the region upstream of *feoABC* and are depicted by superscript 1 (higher transcription rate) and 2 (lower transcription rate). For visual clarity, the relationship between the -10/-35 regions and the BqsR box are also depicted as cartoons, with the orange corresponding to the BqsR box and the black corresponding to the -10/-35 sites. The known regulatory impact of *Pa*BqsR (+ represents induction, - represents repression) on gene expression was based on RNA-seq for cultures grown aerobically in the presence of 4 µM $Fe^{2+}$ or anaerobically in the presence of 100 µM $Fe^{2+}$.[6] For the genes in bold, the transcriptional changes at 4 µM $Fe^{2+}$ have been validated by qPCR. This analysis suggests that the positive mode of *Pa*BqsR regulation correlates with the position of the BqsR box downstream of the -10/-35 regions, whereas all the genes with the negative or the combinatory regulation showed the BqsR box overlapping with at least one of the -10/-35 regions.

**a**

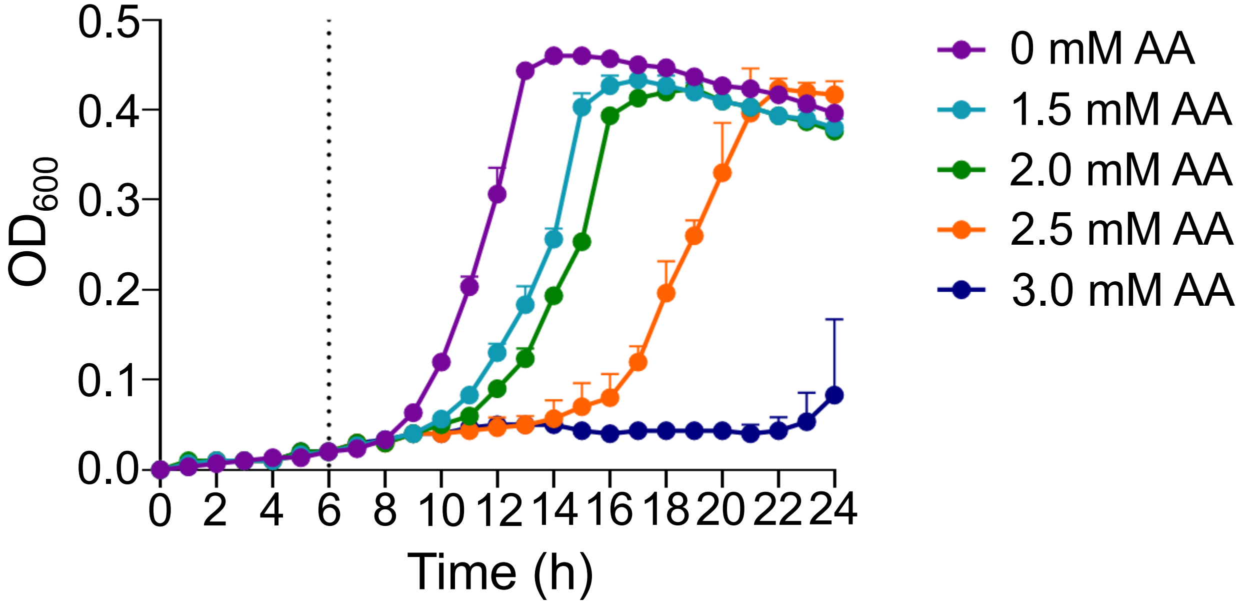


**b**

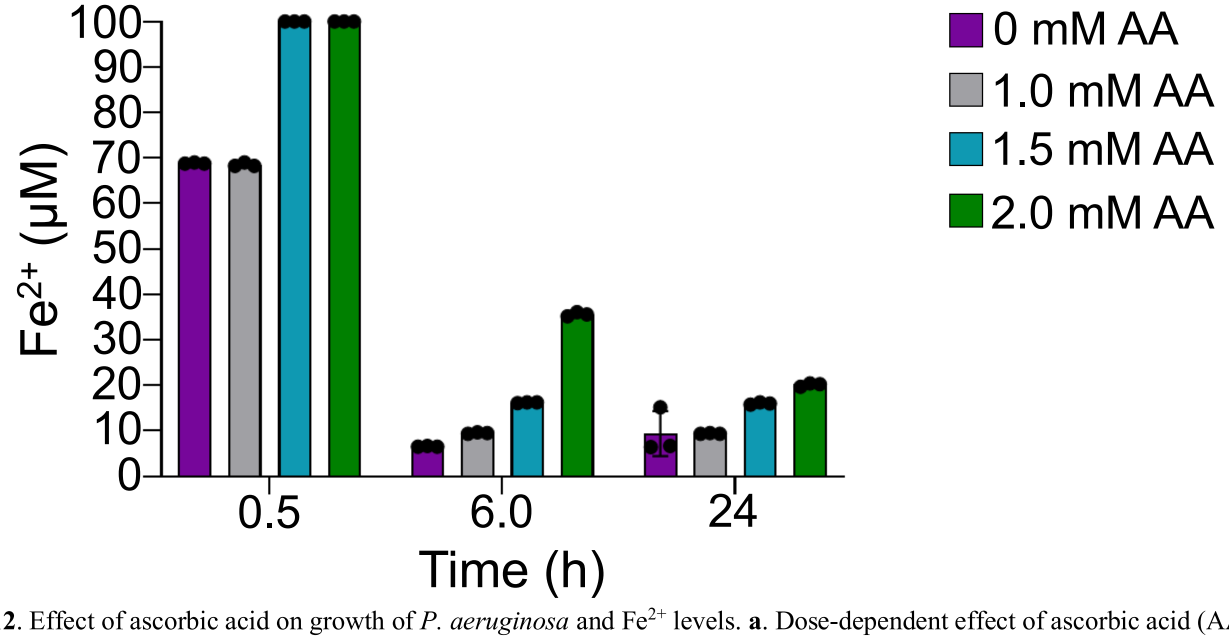


**Figure S12**. Effect of ascorbic acid on growth of *P. aeruginosa* and $Fe^{2+}$ levels. **a**. Dose-dependent effect of ascorbic acid (AA) on growth of WT *P. aeruginosa* grown in BMM8.5 medium with no added $FeSO_4$. Cultures were treated with AA stocks at increasing concentrations or water as a control after six hours of growth (indicated by the vertical dashed line). The optical density at 600 nm was measured every hour for 24 hours. The experiment was performed with three biological replicates for each AA condition. **b**. Dose-dependent effect of AA on maintaining the reduced state of iron ($Fe^{2+}$) over time. BMM8.5 was spiked with 100 μM $FeSO_4$ at increasing concentrations of AA. The concentration of $Fe^{2+}$ was measured after 0.5, 6, and 24 h of static incubation at 37°C using a ferrozine-based assay described in the Methods. The experiment was performed with three independent replicates per condition. Error bars represent ± one standard deviation of the mean.

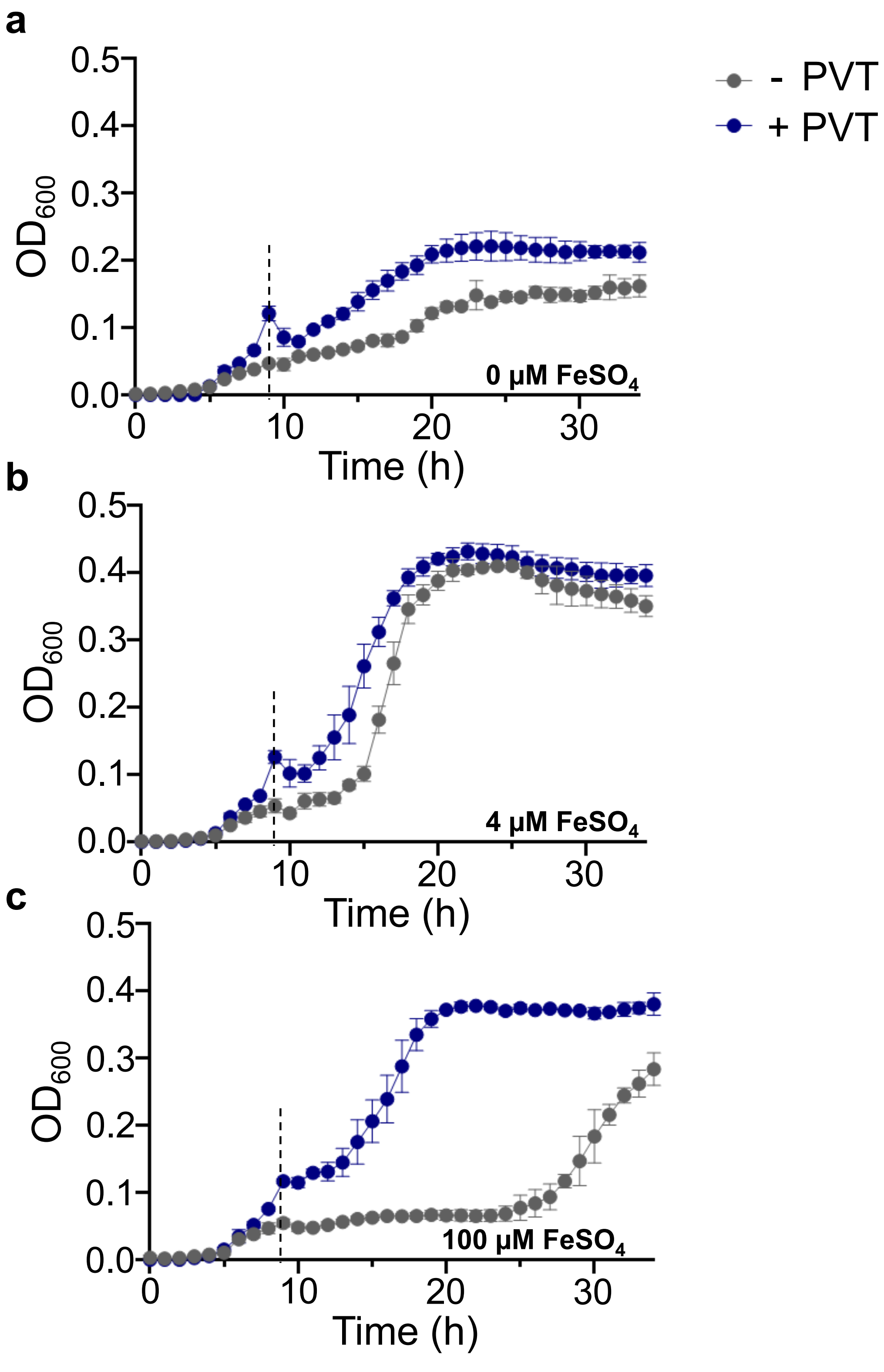


**Figure S13**. Effect of pyruvate (PVT) on *P. aeruginosa* growth at different concentrations of $FeSO_4$. WT *P. aeruginosa* was grown statically at 37 °C in BMM8.5 medium containing 10 mM pyruvate and no iron. After 9 h of growth (indicated by a dashed line), cultures were treated with a final concentration of 0 (**a**), 4 (**b**), or 100 (**c**) µM $FeSO_4$ that had been dissolved in aqueous ascorbic acid for a final concentration of 2 mM in the cultures. The cultures were grown statically at 37 °C for an additional 24 h while the optical densities at 600 nm were measured hourly. The experiment was performed with three biological replicates per strain. Error bars represent ± one standard deviation of the mean.

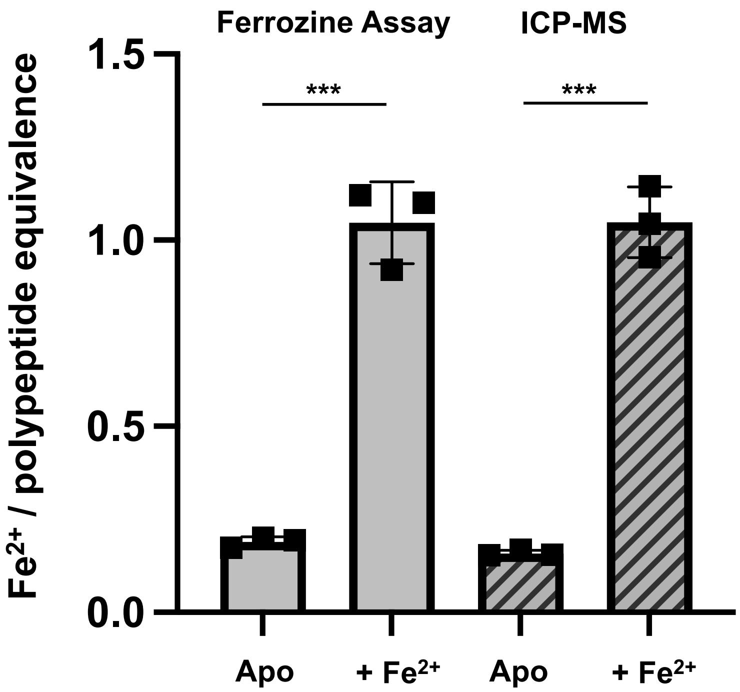


**Figure S14**. Intact, WT *Pa*BqsR binds a single $Fe^{2+}$ ion. Ferrozine assays (*left columns*; $N = 3$) confirm the binding of *ca.* 1 mol. eq. of $Fe^{2+}$ per polypeptide that is confirmed by inductively coupled plasma mass spectrometry (ICP-MS) (*right columns*; $N = 3$). Error bars represent ± one standard deviation of the mean. *** Represents $P < 0.001$ based on the student's t-test.

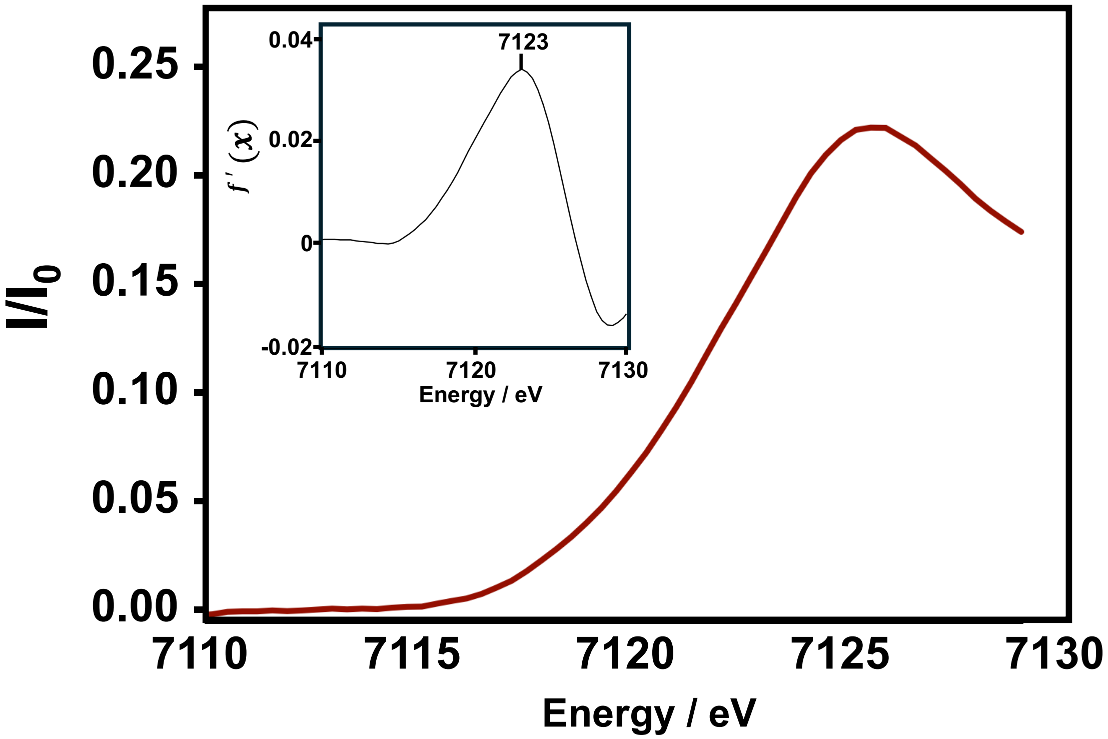


**Figure S15**. X-ray absorption near-edge structure (XANES) spectrum of $Fe^{2+}$-bound WT *Pa*BqsR. *Inset*: First derivative spectrum of the XANES spectrum.

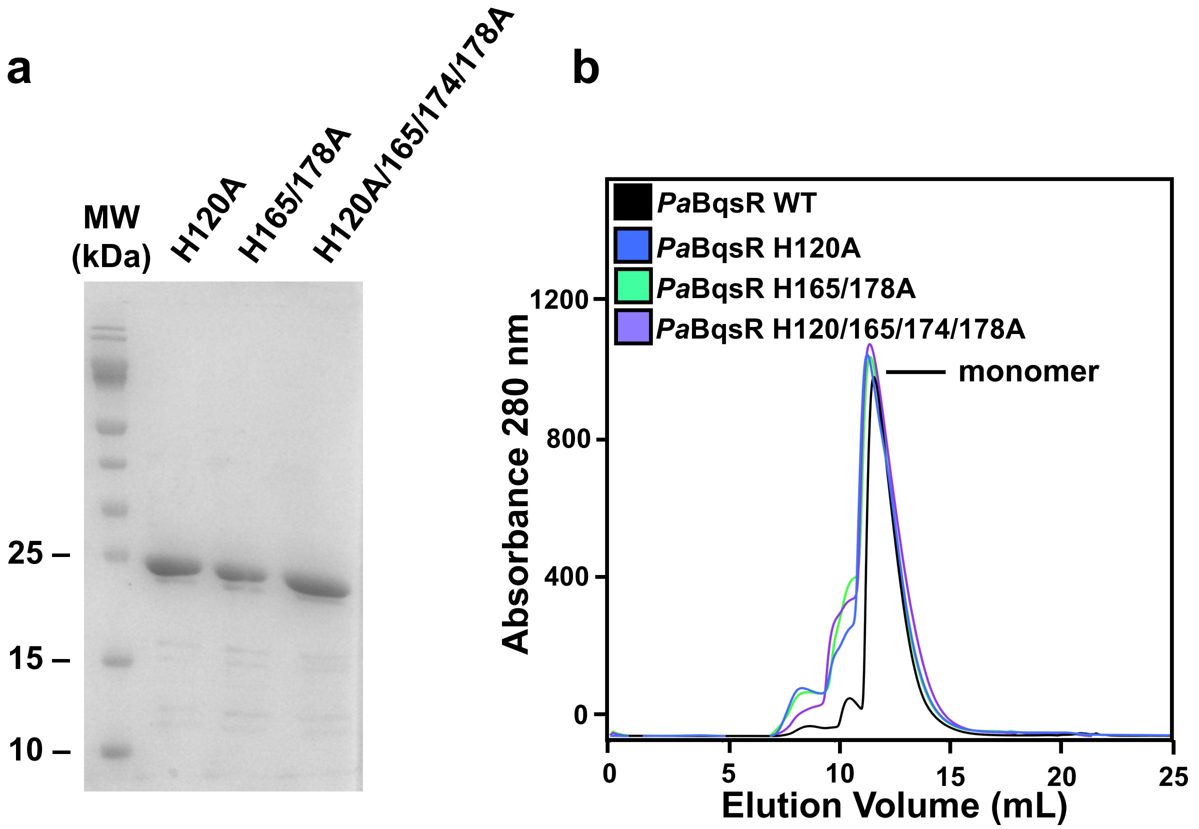


**Figure S16**. Purified *P. aeruginosa* BqsR variants deficient in iron binding. **a.** 15 % SDS-PAGE analysis of *Pa*BqsR variants H120A, H165/178A, and H120/165/174/178A. **b.** WT *Pa*BqsR and *Pa*BqsR variants H120A, H165/178A, and H120/165/174/178A all migrate similarly based on size-exclusion chromatography.
